# Isolation of structural constraints by ensemble-conditioned protein sequence design

**DOI:** 10.1101/2025.09.30.679633

**Authors:** Richard W. Shuai, Ling Zhou, Gina El Nesr, Jessica Lin, Tianyu Lu, Irimpan Matthews, Subhang Bhatti, Justine Yuan, Petr Kouba, Po-Ssu Huang

**Affiliations:** Biophysics Program, Stanford University, Stanford, CA, USA; Department of Bioengineering, Stanford University, Stanford, CA, USA; Stanford Synchrotron Radiation Lightsource, SLAC National Accelerator Laboratory, Menlo Park, CA, USA; Stanford Online High School, Stanford, CA, USA; Independent researcher, San Francisco, CA, USA; CIIRC, Czech Technical University in Prague, Prague, Czech Republic; Loschmidt Laboratories, Masaryk University, Brno, Czech Republic

**Author notes:** Work done as a visiting researcher at Stanford University (Petr Kouba).

## Abstract

Natural sequences have long been the assumed ground truth for protein sequence design methods. But natural sequences encode more than the structural fold, carrying evolutionary history that reflects ancestry, drift, and function. Deep learning models for protein sequence design inherit this signal, achieving higher native sequence recovery but lower predicted foldability. Motivated by this trade-off, we introduce Caliby, a Potts model-based sequence design method capable of conditioning on structural ensembles. By conditioning on synthetic ensembles generated from an input backbone, Caliby samples sequences consistent with shared structural constraints while averaging out native sequence confounders. This strategy improves designability while reducing native sequence recovery, and generalizes across sequence design models. Mechanistically, designed sequences show reduced covariation between non-contacting residues and lower energetic frustration, indicating more precise structural encoding. We demonstrate experimental applicability by designing thermostable proteins, functional lanthanide binders, and soluble membrane protein variants, including a pentameric assembly confirmed by cryo-EM.

## Introduction

The ability to generate amino acid sequences that reliably fold into specified three-dimensional structures is central to protein engineering and *de novo* protein design, with applications from therapeutic biologics to novel enzymes. Deep learning models for structure-conditioned sequence design approach this task by learning structure-sequence rules across large datasets of protein structures, enabling fast and robust design^1–8^.

Because these models are often trained to reproduce native sequences, they can capture features of natural proteins beyond those that specify the fold. Natural sequences arise from evolutionary processes, containing correlations reflecting phylogenetic relatedness, neutral drift, and functional constraints. Additionally, models may inherit biases from the structural datasets on which they are trained. A clue that current models learn non-structural correlations comes from how they behave as they scale: with increasing model capacity, native sequence recovery continues to improve while predicted foldability of designed sequences degrades^6^, suggesting that added capacity is increasingly spent reproducing sequence features beyond what is needed to specify structure.

Broadly, native sequence recovery is a poor proxy for foldability^3,8^. Designs are therefore typically evaluated by their predicted foldability, or designability, which measures how closely a structure prediction model predicts a designed sequence to fold back into the desired structure. However, much of observed protein structure space remains undesignable by current methods^9^, suggesting that isolating structural constraints from these non-structural correlations could expand the space of designable structures.

Energy landscape theory offers a potential explanation for why these non-structural correlations can impair folding. Sequences that fold reliably are thought to have minimally frustrated global energy landscapes^10^, where the native structure sits at the bottom of a smooth funnel. Nevertheless, native proteins are also known to tolerate sites of local energetic frustration that may arise from neutral drift or evolutionary constraints beyond folding, such as function or binding^11,12^. As a result, models trained on native proteins may learn to reproduce these local frustration patterns, introducing conflicting interactions that may hinder the correct folding of designed sequences.

Motivated by these observations, we introduce Caliby, a Potts model-based sequence design method built on the idea that conditioning on synthetic structural ensembles can suppress learned sequence patterns that conflict with folding. Given an input backbone, we first generate a synthetic ensemble by partial diffusion with a backbone generative model^13^, then design sequences with Caliby conditioned on this ensemble (Fig. 1a,b). This yields sequences satisfying structural constraints shared across the ensemble, while averaging out non-structural correlations specific to native proteins (Fig. 1c). The resulting designs more precisely match native backbones under single-sequence AlphaFold2^14^, with lower native sequence recovery (Fig. 1d), an effect that transfers across model families and coincides with both suppressed non-structural sequence covariation and reduced local energetic frustration^11^. Finally, we experimentally characterize designs spanning thermostable proteins, functional lanthanide binders, and soluble membrane protein variants, including a pentameric assembly confirmed by cryo-EM.

**Figure 1.**
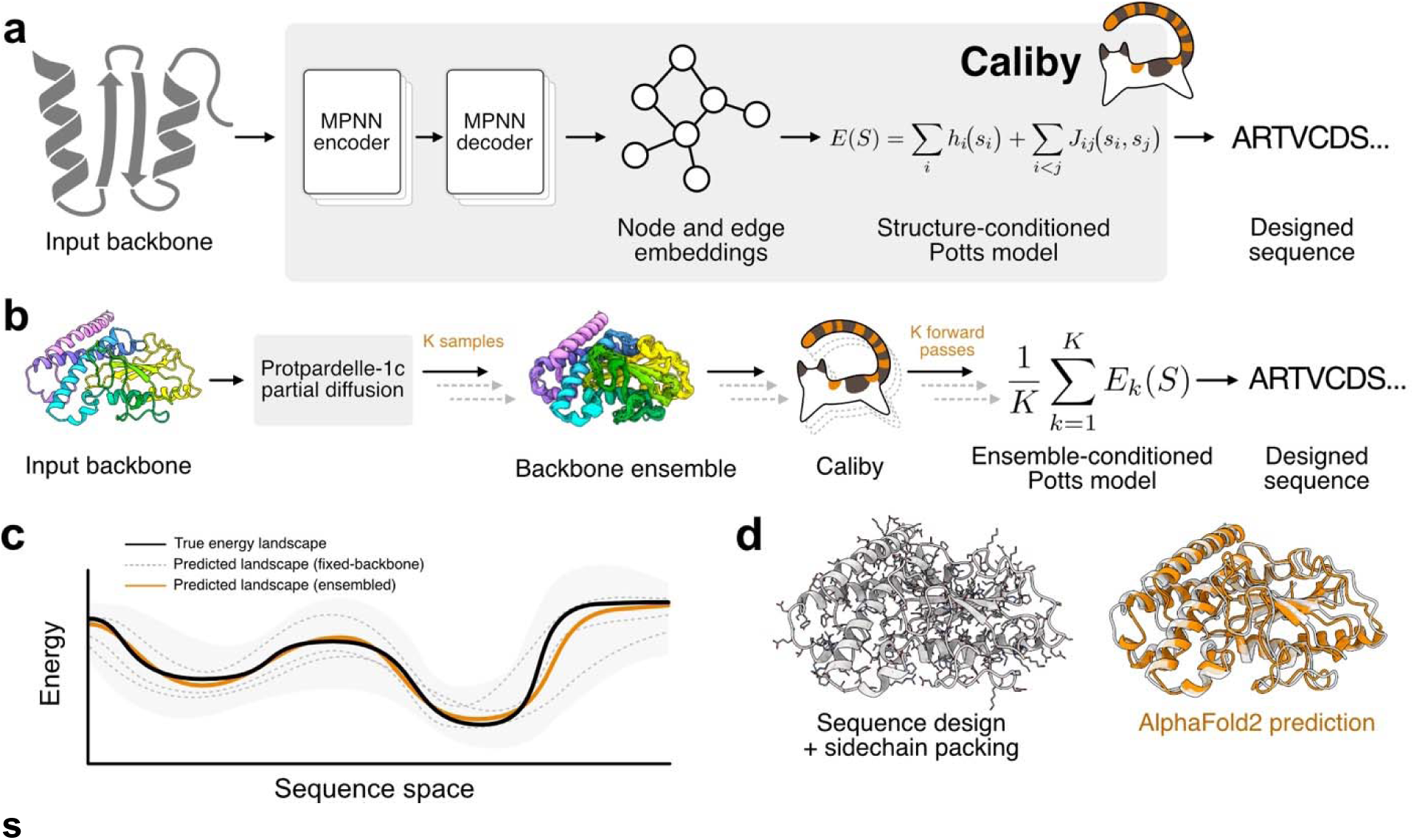
Caliby overview. **(a)** In fixed-backbone mode, Caliby designs sequences by sampling from a structure-conditioned Potts model derived from a graph neural network. **(b)** Caliby conditions on a structural ensemble by averaging the sitewise and pairwise energy terms of per-structure Potts models, defining the energy of a sequence as its mean energy across the ensemble. **(c)** Each backbone in a structural ensemble yields a Potts model that defines a predicted energy landscape over sequences (gray dashed lines; shaded region shows spread). These landscapes vary across backbones but tend to agree at points reflecting shared structural constraints. Averaging across the ensemble produces a predicted landscape (orange) that more closely matches the true energy landscape (black). **(d)** The full design pipeline performs sequence design with Caliby and sidechain packing with CalibyPacker (left), yielding sequences whose single-sequence AlphaFold2 predictions (right, orange) closely match structurally complex target backbones (right, gray).

## Results

### Caliby designs sequences from structural ensembles

Caliby builds upon the ProteinMPNN architecture but replaces the autoregressive sequence decoding with prediction of Potts model parameters, similar to other Potts-based design methods^4,8,15^ (Methods). We removed the causal mask from the decoder layers and adapted the ChromaDesign Potts decoder module, which derives sitewise and pairwise energy terms from node and edge embeddings^4^. Following ChromaDesign, Caliby samples sequences from the resulting neural-network-derived Potts model by Discrete Langevin Monte Carlo (DLMC)^16^ with a local composition perplexity (LCP) restraint to penalize unnaturally low complexity sequences.

Although Caliby was trained on static structures from the Protein Data Bank (PDB)^17^, the Potts model formulation provides a straightforward mechanism for conditioning on structural ensembles. To condition on an ensemble of structures, we define the energy of a sequence as the average of its energies under each per-structure Potts model, which is equivalent to scoring under a single Potts model with energy terms averaged over the ensemble. Once this averaged model is precomputed, sampling costs no more than for a single structure. Because the resulting sequence distribution is the normalized geometric mean of the per-structure distributions, related to a Product-of-Experts model^18^, ensemble-conditioned design samples from the distribution of sequences compatible with all structures in the ensemble (Supplementary Note 1).

To generate synthetic ensembles from an input backbone, we used Protpardelle-1c^13^ partial diffusion with 150 rewind steps out of 500 total steps to generate 31 additional backbone conformers (K = 32). We found that (K = 32) provides strong results, with slightly improved but diminishing returns for larger ensemble sizes (Extended Data Fig. 1). Our approach generalizes to other conformer generation methods: ensembles generated by RFdiffusion319 partial diffusion performed comparably to Protpardelle-1c, whereas ensembles generated by adding Gaussian noise tolerated less structural variation before degrading (Extended Data Fig. 1).

To complete the design pipeline, we also trained a dedicated model (CalibyPacker) for sidechain packing using a diffusion-based approach similar to FAMPNN^7^ (Fig. 1d, Methods). CalibyPacker achieves competitive sidechain packing accuracy relative to prior methods, as measured by performance on a set of CASP13, CASP14, and CASP15 targets held out at 40% sequence similarity following prior benchmarks^7,20^ (Supplementary Fig. 1).

### Ensemble conditioning improves designability while reducing native sequence recovery

If ensemble conditioning isolates structural constraints, designed sequences should be predicted to fold more reliably to their target backbones. We tested this hypothesis on a held-out set of native backbones of lengths 32 to 512, sampling 8 sequences per backbone with fixed-backbone Caliby (Caliby-fb), ensemble-conditioned Caliby with 32 conformers (Caliby-e32), ProteinMPNN, and ChromaDesign. We then assessed the best-of-8 by single-sequence AlphaFold2 pLDDT and self-consistency TM score (scTM). As predicted, Caliby-e32 achieved the best self-consistency on native backbones, enabling the design of challenging structures with previously low designability (Fig. 2a, Extended Data Fig. 2).

**Figure 2.**
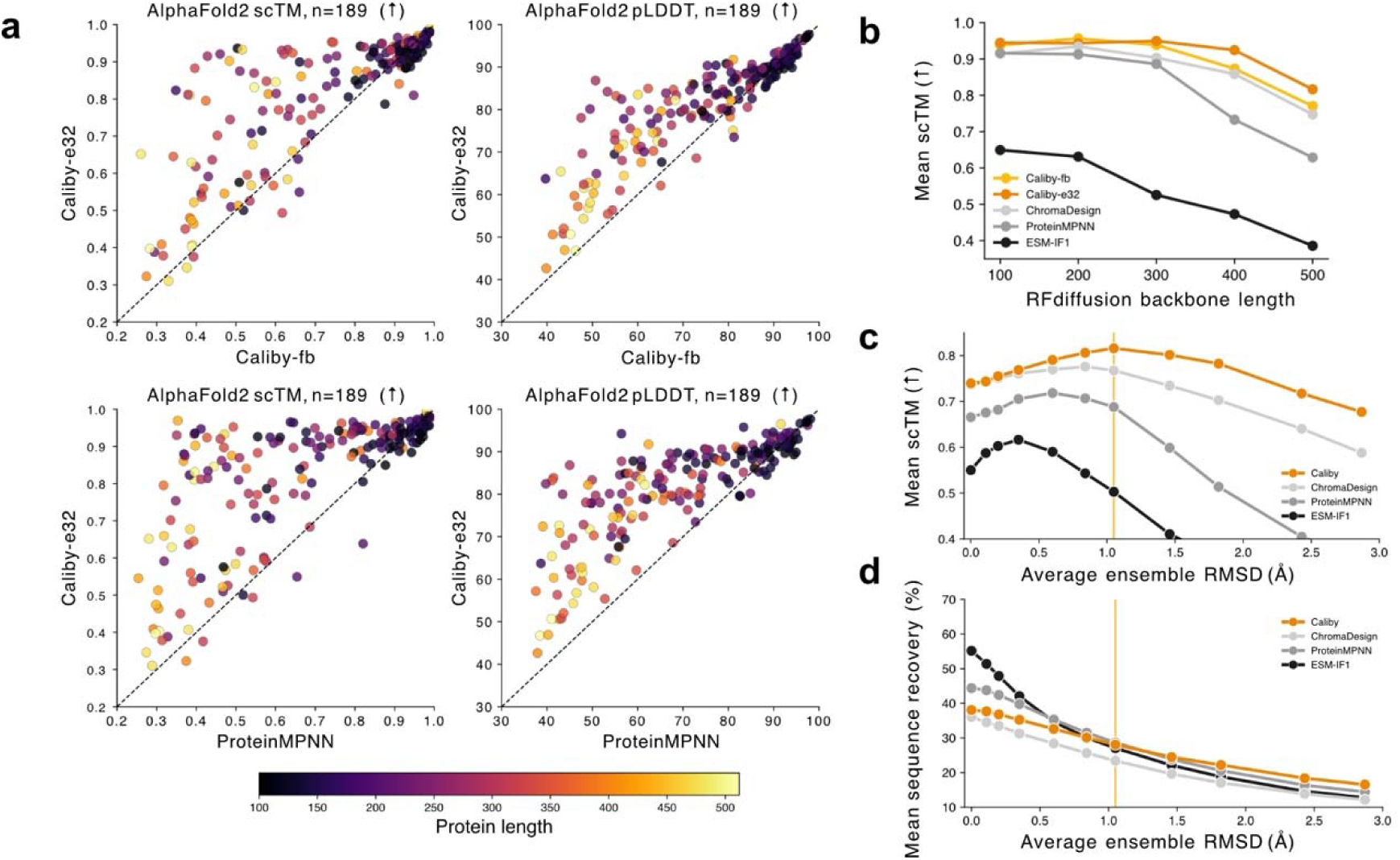
Ensemble conditioning improves designability. **(a)** Caliby conditioned on a synthetic ensemble of 32 conformers (Caliby-e32) outperforms fixed-backbone Caliby (Caliby-fb, top) and ProteinMPNN (bottom) based on single-sequence AlphaFold2 self-consistency (scTM, left; pLDDT, right) across n = 189 native backbones of lengths 32-512. Best of 8 designs shown per backbone, colored by protein length. **(b)** Mean scTM as a function of RFdiffusion backbone length for Caliby-fb, Caliby-e32, ChromaDesign, ProteinMPNN, and ESM-IF1 (n = 100 backbones per length). **(c)** Mean scTM over n = 189 native backbones as a function of ensemble structural diversity for Caliby, ChromaDesign, ProteinMPNN, and ESM-IF1. Ensemble conditioning is implemented by Potts energy averaging for Caliby and ChromaDesign and by tied sampling for ProteinMPNN and ESM-IF1 with 32 conformers. 0Å represents fixed-backbone design. The orange line represents 150 Protpardelle-1c rewind steps, used as the Caliby default. **(d)** Mean native sequence recovery across the same n = 189 native backbones as a function of ensemble diversity.

Ensemble conditioning also improved designability on *de novo* backbones from RFdiffusion and Protpardelle-1c, particularly for longer, more complex backbones (Fig. 2b, Supplementary Fig. 2). Caliby’s LCP restraint, adopted from ChromaDesign, helps to avoid repetitive, low-complexity sequences designed by prior methods (Extended Data Fig. 3). Additionally, ensemble-conditioned Caliby improved designability over SolubleMPNN^21^ for binders designed by Protpardelle-1c, especially on binder-only self-consistency metrics, but not for BindCraft-designed binders^22^, likely because BindCraft produces highly idealized backbones designable by either method (Extended Data Fig. 4).

If averaging across synthetic ensembles isolates structural constraints, the effect should not be specific to Caliby. We therefore applied ensemble conditioning across model families. For the Potts-based methods, Caliby and ChromaDesign, we averaged Potts parameters across the ensemble. For the autoregressive methods, ProteinMPNN and ESM-IF1^2^, we used tied sampling. For each method, we swept ensemble structural diversity by tuning the number of Protpardelle-1c rewind steps used to generate conformers, quantifying diversity as the average RMSD of the synthetic backbones to the starting structure. In every case, designability improved while native sequence recovery declined, demonstrating that this behavior is not model-specific (Fig. 2c, 2d). Past an optimal point, ensembles become too structurally diverse to be beneficial for sequence design. Overall, Caliby performs best for designability and tolerates the greatest ensemble diversity before degrading. Furthermore, relative to tied sampling methods, Potts parameter averaging scales more computationally efficiently with ensemble size and defines the ensemble-conditioned distribution in closed form rather than approximating it stepwise (Supplementary Note 1).

To support the hypothesis that larger models learn to capture sequence patterns in native sequences even when these patterns do not strongly correlate with structure, we also trained Caliby-L, a larger version of Caliby where we increased the number of MPNN layers from 3 to 5 and increased the hidden dimension from 128 to 256. Caliby-L achieves higher test set sequence recovery but lower scTM than Caliby, suggesting that the increased model capacity captures native sequence biases unrelated to foldability (Extended Data Fig. 5). However, by using ensemble conditioning with Caliby-L, we improved the designability of sequences to levels close to that of Caliby while similarly reducing native sequence recovery. This provides additional evidence that ensemble-conditioned design can average out the non-structural, native sequence biases learned by larger models.

While native sequence recovery drops with increasing ensemble diversity, consensus sequence recovery and similarity to MSA-derived PSSMs decline more slowly, suggesting that ensemble conditioning suppresses idiosyncratic sequence features (Extended Data Fig. 1). Beyond improving designability, ensemble-conditioned Caliby increases the positional entropy of designed sequences both on the protein surface and the core (Extended Data Fig. 5). Ablations on ensemble size, sampling temperature, and conformer generation support our default sampling settings (Extended Data Fig. 1). Notably, designing on a single synthetic conformer did not reproduce the improvement, demonstrating that the performance gain is not attributable to backbone refinement alone (Extended Data Fig. 1c).

### Ensemble conditioning suppresses learned correlations unrelated to structure

We hypothesized that ensemble-conditioned design improves single-sequence AlphaFold2 designability by suppressing learned spurious correlations unrelated to structural constraints. To provide evidence for this, we performed direct-coupling analysis (DCA) to predict structural contacts^23^ on both natural multiple sequence alignments (MSAs) and synthetic MSAs designed by Caliby at equal MSA depths (Fig. 3a). Because DCA relies on observed correlations in the MSA for contact prediction, DCA struggles to disentangle true structural contacts from correlations induced by other constraints, including protein function and phylogeny^24^. Sweeping sampling temperature to control for MSA diversity, both Caliby-fb and Caliby-e32 reached higher peak long-range contact precision (P@L) than natural MSAs, and Caliby-e32 exceeded Caliby-fb, suggesting that sequences designed with ensemble conditioning tend to have reduced covariation between non-contacting residue pairs.

**Figure 3.**
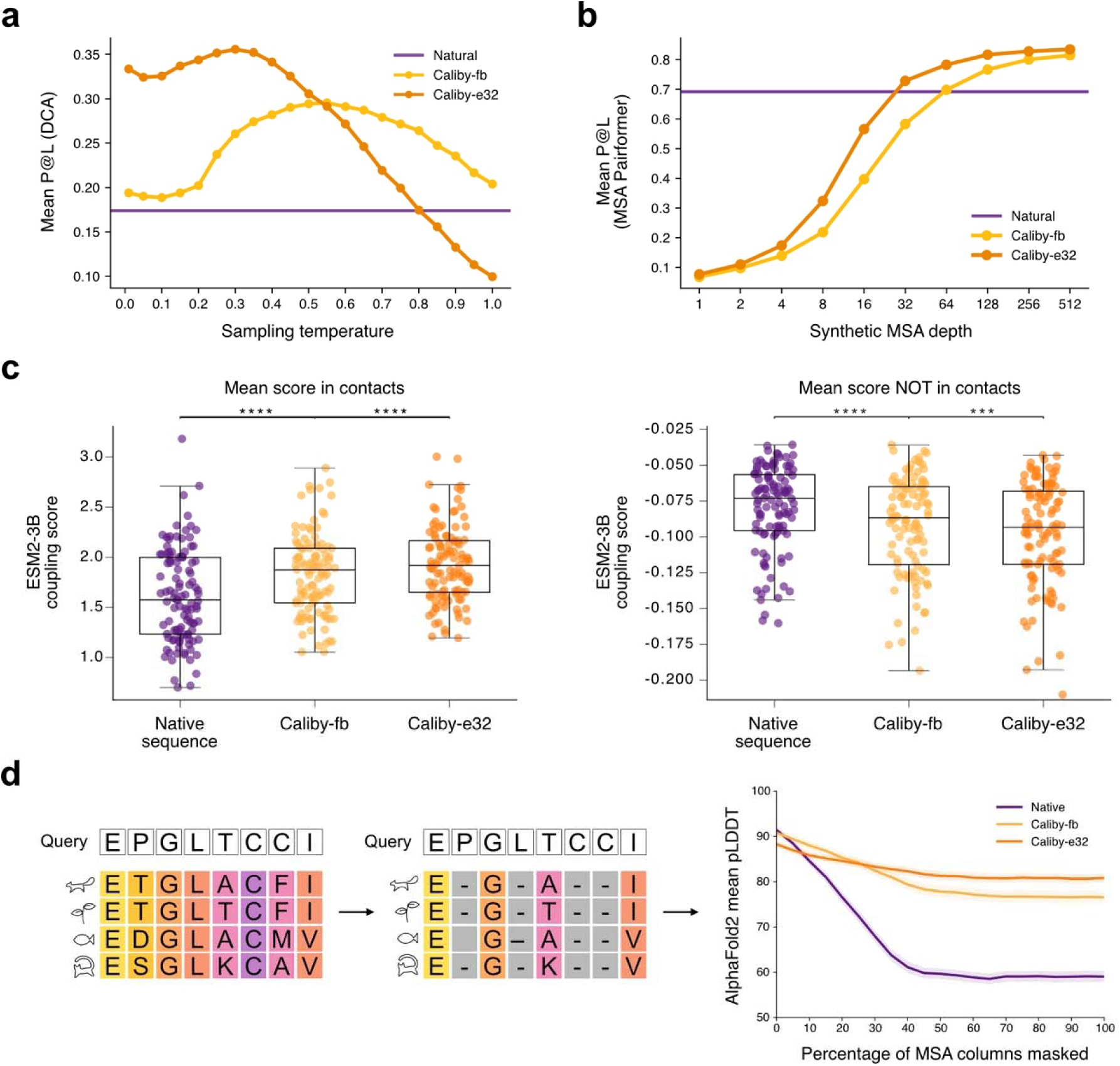
Ensemble conditioning suppresses sequence covariation unrelated to structure. **(a)** Long-range contact precision over the top L residue pairs (P@L) from direct coupling analysis on natural MSAs (purple) and on synthetic MSAs from Caliby-fb and Caliby-e32 across sampling temperatures (n = 96). **(b)** P@L from MSA Pairformer as a function of synthetic MSA depth, compared with a natural sequence alignment of 512 sequences (purple) (n = 96 structures). **(c)** ESM-2 3B categorical Jacobian magnitudes averaged over contacting (left) and non-contacting (right) residue pairs for native sequences, Caliby-fb designs, and Caliby-e32 designs with n = 111 structures (***P < 0.001, ****P < 0.0001, two-sided Wilcoxon signed-rank test). **(d)** Mean AlphaFold2 pLDDT as MSA columns are progressively masked with gap tokens, interpolating from full coevolutionary information (0%) to single-sequence prediction (100%) (n = 111 structures, 8 designed sequences per structure, 3 random seeds per masking level). Shaded regions show ± 1 SEM.

We performed a similar contact prediction analysis with MSA Pairformer^25^ by predicting contacts for a query sequence given an MSA (Fig. 3b). Synthetic MSAs reached the contact precision of a 512-sequence natural alignment at much shallower depths, with roughly 32 sequences for Caliby-e32 and 64 for Caliby-fb, paralleling the DCA results.

As a complementary analysis using a large protein language model, we computed categorical Jacobians^26^ for native and designed sequences with the ESM-2 3-billion parameter model^27^ (Fig. 3c). Coupling strength at structural contacts increased from native to Caliby-fb to Caliby-e32 sequences, while couplings at non-contacting residue pairs decreased. Finally, we assessed this hypothesis directly with AlphaFold2 We interpolated between using full co-evolutionary information and using the query sequence by predicting structures with the full MSA and then progressively masking MSA columns (Fig. 3d). We found that sequences from ensemble-conditioned Caliby require less coevolutionary information for accurate prediction. These results support the hypothesis that ensemble conditioning suppresses learned correlations unrelated to structural constraints, yielding sequences whose covariation more precisely reflects backbone geometry.

### Ensemble-conditioned Caliby reduces local energetic frustration

Energy landscape theory predicts that reliably folding sequences have minimally frustrated energy landscapes^10^, but native proteins tolerate local frustration arising from evolutionary constraints unrelated to folding^11^. We hypothesized that sequence design models may spuriously reproduce these patterns, introducing conflicting interactions that hinder correct folding. To test whether ensemble conditioning mitigates this, we used Frustratometer^11,12^ to compute local frustration indices under the Associative Memory, Water Mediated, Structure and Energy Model (AWSEM) energy function^28^. We computed frustration along an interpolation path that progressively biases Caliby’s sampling toward the native sequence using a parameter λ, ranging from full redesign at λ = 0.0 to the native sequence at higher λ (Methods).

For both Caliby-fb and Caliby-e32, decreasing λ yields lower sequence recovery and higher scTM (Fig. 4a). Along these interpolation paths, average frustration decreases as Caliby redesigns more of the native sequence, and Caliby-e32 reduces frustration more than Caliby-fb (Fig. 4b, top, Extended Data Fig. 6), suggesting that ensemble conditioning promotes smoother energy landscapes. Applying standard frustration thresholds^11^, Caliby-e32 also reduces the percentage of highly frustrated sites and increases the percentage of minimally frustrated sites relative to the native sequence (Fig. 4b, bottom).

**Figure 4.**
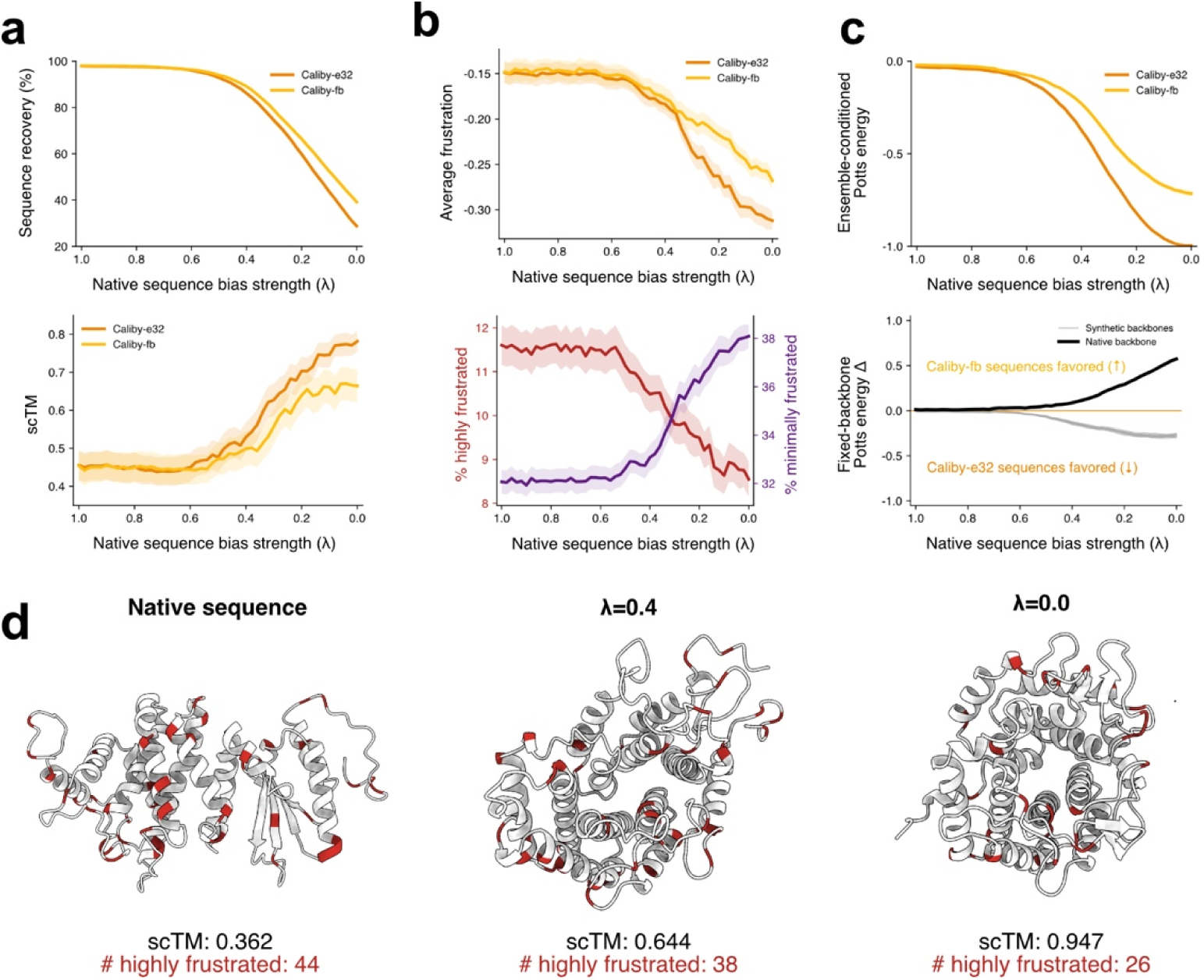
Ensemble conditioning reduces local energetic frustration. **(a)** Sequence recovery (top) and scTM (bottom) as a function of native sequence bias strength λ, ranging from strongest native bias (λ = 1) to full redesign (λ = 0) (n = 27 backbones, 8 designed sequences per backbone at each). (b) Average frustration (top), defined as the negated single-residue mutational frustration index from Frustratometer, for Caliby-fb and Caliby-e32. Percentage of highly frustrated and minimally frustrated residues for Caliby-e32 (bottom). **(c)** Ensemble-conditioned Potts energy (top) and fixed-backbone Potts energy difference (Caliby-e32 minus Caliby-fb; bottom) along the interpolation path (n = 27 backbones). Energies are normalized per backbone such that the native sequence scores 0 and the lowest-energy sequence across both interpolation paths scores -1, then averaged across backbones. In the bottom panel, positive values indicate Caliby-fb sequences are more favorable. Caliby-fb sequences score more favorably on the native backbone (black) but worse on synthetic conformers (gray) relative to Caliby-e32. **(d)** Example backbone illustrating the reduction in highly frustrated sites (red) and improvement in scTM as λ decreases. Shaded regions in (a-c) show ± 1 SEM.

Scoring sequences along the interpolation paths with Caliby’s Potts model reveals that fixed-backbone design over-optimizes for a single conformation (Fig. 4c). Under the ensemble-conditioned energy, both Caliby-fb and Caliby-e32 reach lower energies than native-like sequences (Fig. 4c, top). However, scoring against individual structures separates them: relative to Caliby-e32, Caliby-fb sequences are favored on the native backbone but disfavored on the synthetic conformers, indicating narrow optimization to a single conformation rather than the surrounding ensemble (Fig. 4c, bottom). This parallels the reduced local frustration observed with ensemble-conditioned designs. We illustrate this reduction with an example backbone, where the number of highly frustrated sites decreases and scTM increases as design becomes less constrained (Fig. 4d).

### Caliby designs thermostable, soluble, and functional proteins

Although designability metrics correlate with experimental success^22,29^, they do not guarantee experimental viability. We therefore experimentally characterized Caliby designs, first by redesigning native backbones, then by engineering a functional metal-binding site.

For native backbone redesign, we selected 32 monomeric, single-domain apo structures from the test set. Similar to SolubleMPNN^21^, we trained SolubleCaliby, a variant of Caliby trained by excluding transmembrane proteins from the training set (Methods). We compared SolubleCaliby-e32 (16 samples) and SolubleMPNN (512 samples), selecting the best sequence by scTM. Despite using 32-fold fewer samples, SolubleCaliby-e32 surpassed SolubleMPNN’s best-of-512 in scTM, demonstrating that ensemble conditioning achieves high designability with substantially less sampling (Fig. 5a, top). From these 32 backbones, we selected 12 diverse backbones spanning the full range of pLDDTs achievable by SolubleMPNN and experimentally characterized designs from both methods (Fig. 5a, bottom; Supplementary Tables 3 and 4). Two were transmembrane proteins, requiring the design of soluble variants of membrane-associated folds.

**Figure 5.**
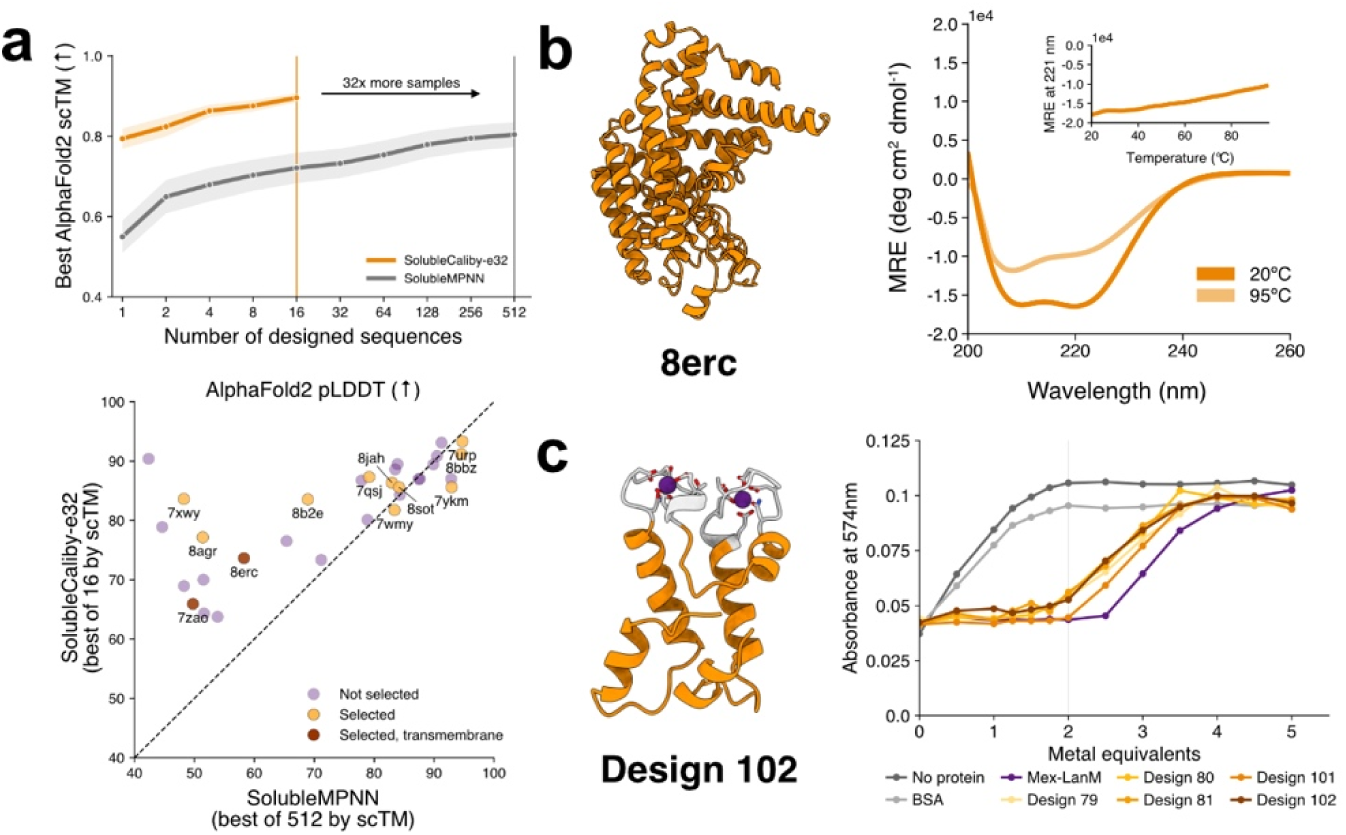
Caliby designs thermostable, soluble, and functional proteins. **(a)** Top, best-so-far single-sequence AlphaFold2 scTM as a function of the number of designed sequences for SolubleCaliby-e32 and SolubleMPNN across n = 32 monomeric, single-domain apo backbones from the test set. The orange vertical line denotes 16 sequences, and shaded regions denote ± 1 SEM. Bottom, AlphaFold2 pLDDT for the best-by-scTM SolubleCaliby-e32 design of 16 and best-by-scTM SolubleMPNN design of 512. The 12 backbones selected for experimental characterization are labeled by PDB code. **(b)** AlphaFold2 prediction of the SolubleCaliby-e32 design for membrane-bound O-acyltransferase 8ERC (left) with CD wavelength scans at 20°C and 95°C (right). Thermal denaturation was monitored by mean residue ellipticity at 221 nm (inset). **(c)** Boltz-2 prediction of SolubleCaliby-e32 design 102 (left), scaffolding the lanthanide-coordinating EF-hand loops of *Hans*-LanM (gray, fixed motif). Xylenol orange competition assay (right) reporting free Sm^3+^ by absorbance at 574 nm as a function of metal equivalents, for xylenol orange alone, BSA as a non-binding control, the native lanmodulin Mex-LanM, and 5 designs. The gray vertical line marks two equivalents, which is the expected stoichiometry from the two fixed EF-hand loops per design.

Eleven of 12 SolubleCaliby designs and 10 of 12 SolubleMPNN designs yielded soluble, folded proteins as assessed by size exclusion chromatography (SEC) and circular dichroism (CD) (Extended Data Fig. 7, 8). The SolubleMPNN design for a membrane-bound O-acyltransferase (PDB: 8ERC)^30^ was found predominantly in the insoluble fraction (Supplementary Fig. 3), whereas the corresponding SolubleCaliby design eluted as a monodisperse peak with CD spectra and thermal denaturation consistent with a well-folded, thermostable protein (Fig. 5b). Both methods failed on the backbone of membrane-associated adhesin from *Brachyspira pilosicoli* (PDB: 7ZAO)^31^, producing proteins that lacked well-defined secondary structure by CD (Extended Data Fig. 8).

Many design applications require scaffolding a fixed functional motif rather than redesigning an entire backbone. To test whether Caliby can support precise geometry at a binding site, we designed lanthanide-binding proteins, which are of interest for selective rare-earth recovery^32^. We used SolubleCaliby-e32 to design non-motif positions of Protpardelle-1c backbones scaffolding the lanthanide-coordinating EF-hand loops of the lanmodulin from *Hansschlegelia quercus* (*Hans*-LanM; PDB: 8DQ2)^32^ and characterized 5 designs (Methods). We assessed metal binding using a competition assay with xylenol orange (XO), a colorimetric indicator that reports free Sm^3+^ (Fig. 5c). Whereas XO alone and a non-binding control BSA reported free metal immediately, all 5 designs suppressed the XO signal until approximately two equivalents of Sm^3+^. This indicates that each design sequesters close to two metal ions, as expected from the two fixed EF-hand loops in each design. A native lanmodulin, Mex-LanM, suppressed the signal to higher metal equivalents, reflecting its greater number of coordinating sites^33^.

### Caliby designs soluble membrane protein assemblies from predicted structures

We next applied Caliby to design eight homo-oligomeric membrane protein assemblies from QSProteome^34^, a repository of AlphaFold-predicted quaternary structures, spanning 3-to 10-fold symmetry. In each case, sequence design was conditioned on a predicted model rather than an experimental structure. We designed sequences with SolubleCaliby-e32 for each assembly, tying equivalent positions across chains to enforce symmetry, then filtered designs by both AlphaFold2 and AlphaFold-Multimer^35^ self-consistency (Methods). Four of the eight targets passed, and we chose one design from each for experimental characterization (Extended Data Fig. 9). Of these four designs, three successfully expressed at the 500-mL scale in *E. coli.* At 10 nM, mass photometry droplet-dilution measurements showed that two designs formed a single dominant species corresponding to the expected oligomeric state and were therefore prioritized for cryo-EM screening (Extended Data Fig. 9b-d).

We determined the cryo-EM structure of the design for Q8XAZ3 (YneE), annotated as a voltage-dependent anion channel-forming protein, for which no experimental structure has been reported. The resulting 3.5 Å resolution reconstruction (Extended Data Fig. 10) displayed well-resolved secondary structure elements and sidechain density sufficient for atomic model building (Fig. 6a,b; Supplementary Table 1). The experimentally determined cryo-EM structure closely matched the design input, with a Cα RMSD of 1.06 Å across all 214 atom pairs in each protomer (Fig. 6c). Structural differences were largely confined to surface-exposed loops, consistent with their expected conformational flexibility. The pentameric assembly also exhibited extensive interprotomer packing interactions, confirming that Caliby accurately designed both the tertiary structure of the protomer and the quaternary architecture of the complex. Together, these results show that Caliby can design soluble variants of homo-oligomeric membrane proteins using only AlphaFold-predicted structural models, expanding the range of membrane proteins amenable to computational design.

**Figure 6.**
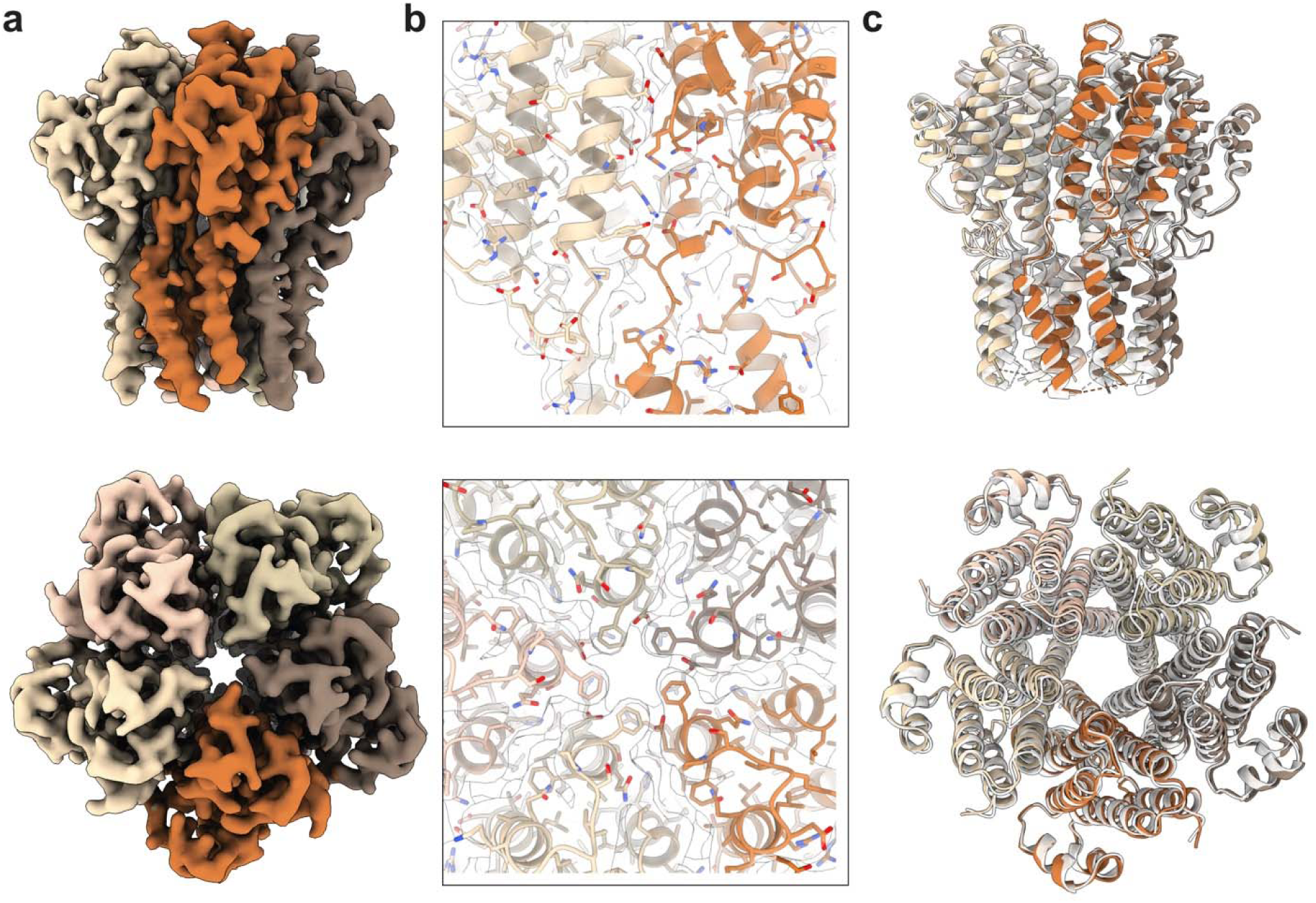
Cryo-EM structure of a Caliby-designed soluble variant of the voltage-dependent anion channel-forming protein YneE (UniProt: Q8XAZ3). (a) A 3.5 Å cryo-EM reconstruction of the Caliby-designed YneE pentamer. (b) Two representative views of the cryo-EM density map (transparent contour) overlaid with the atomic model show well-resolved secondary structure elements and bulky side chains, enabling model building of the pentameric assembly. (c) Following the exclusion of flexible regions that were not resolved in the cryo-EM map, the experimentally determined structure (colored ribbons) closely matches the AlphaFold3-predicted model of the Caliby-designed sequence (white ribbons), demonstrating high structural agreement.

## Discussion

Ensemble-conditioned Caliby enables sequence design for structurally complex backbones, validated here for the design of thermostable soluble proteins, functional lanthanide binders, and a homo-oligomeric membrane protein assembly solved by cryo-EM. Concurrent work has also successfully used Caliby to design the scaffold of a *de novo* bi-zinc metalloenzyme^36^, with the designed structure confirmed by crystallography. In a separate effort, Caliby was used by an autonomous language model agent for de novo binder design, yielding experimentally validated binders across multiple targets^37^. Collectively, these results establish Caliby as a broadly useful tool for structure-conditioned sequence design for diverse functional tasks.

Prior work has shown that native sequence recovery and designability are often not well correlated^3,6,8^. Our work indicates that this gap arises because models trained on native proteins learn to recover phylogenetic correlations, functional constraints, and other sequence biases that conflict with robust folding. We show that averaging over ensembles suppresses these biases, yielding sequences with reduced local energetic frustration^11,12^, consistent with the smooth, funneled energy landscapes theorized to enable robust folding^10^.

Ensemble averaging improves designability across diverse model architectures, suggesting that this behavior is inherent to training on native protein data. A similar trend has been observed in protein language models for fitness prediction, where larger models achieve lower perplexity scores yet perform worse on variant effect prediction^38^, until averaging strategies suppress phylogenetic noise^39^. This reinforces the view that disentanglement of evolutionary signal may be necessary for broadly generalizable protein models.

Finally, the principles underlying our ensemble-conditioning approach are not limited to synthetic ensembles and can be extended to any user-provided ensemble. Therefore, beyond improving designability and enabling functional design, we anticipate that Caliby may be used for robust multistate design^40–42^, where sequences must be compatible across multiple functionally relevant conformations.

## Methods

### Model architecture and training

#### Caliby Architecture

Caliby uses a graph neural network encoder-decoder that builds on the message-passing architecture of ProteinMPNN, with all hidden dimensions set to 128. A k-nearest-neighbor graph (K = 48) is constructed over Cα distances, and edge features encoding pairwise interatomic distances and relative residue positions are computed following ProteinMPNN. The network consists of 3 encoder layers followed by 3 decoder layers. Similar to Full-Atom MPNN (FAMPNN), we removed the causal mask from the decoder layers and expanded the featurization to encode pairwise distances between sidechain atoms if sidechains are provided as context.

After the final decoder layer, the node embeddings and edge embeddings are decoded into the sitewise and pairwise parameters h and j of a pairwise Markov random field (Potts model) over amino acid identities, following the approach from ChromaDesign. Sitewise fields are computed as 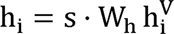, where is a learned global scale initialized at 0.1. Pairwise couplings are parameterized as a matrix product J_ij_ = s · 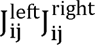, where each factor is linearly projected from the edge embeddings. Both h and j are placed in zero-sum gauge, and j is symmetrized by averaging *j_ij_* with 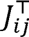. The Potts energy of a sequence S is computed as 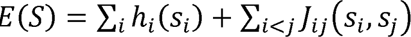, where lower energies are more favorable.

#### Sequence sampling

Following ChromaDesign, sequences are sampled from the Potts model using Discrete Langevin Monte Carlo (DLMC), which updates all positions in parallel at each sweep. We use 500 sweeps with linear temperature annealing from T = 1.0 to T = 0.01 over the first 80% of sweeps. Throughout this work, when sampling temperature is varied, the final annealed temperature is adjusted while keeping the initial temperature fixed at T = 1.0.

To discourage low-complexity solutions, during sampling, we apply the local composition perplexity (LCP) restraint from ChromaDesign. This restraint applies a quadratic penalty on low-entropy local windows:

Where S_i_ is the entropy of window *i*, 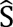 is a threshold entropy value, and 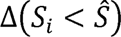 is an indicator of whether S*_i_* is below 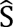. As in ChromaDesign, we set W = 30 and 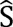 to the 5^th^ percentile of the 30-residue local window entropy in PDB sequences (2.32 nats).

#### Caliby training details

Caliby was trained on protein chains from PDB biological assemblies through the AtomWorks framework, using structures released before 2021-09-30. During training, following prior work, Gaussian noise with standard deviation 0.3Å was added to the input atom coordinates. Following FAMPNN^7^, we also provided partial sequence and sidechain context to the model during training so that users can condition on mixtures of sequence-only and sequence-and-sidechain context. Specifically, we sampled p_1_, p_2 ∽_ Uniform(0,1) for each training example and unmasked each residue identity in the native sequence with probability p_1_. For each residue whose identity was unmasked, we unmasked the native sidechain conformation with probability p_2_. Similar to other work^4,8,15^, we trained Caliby to condition on an input structure and predict a set of Potts parameters over the sequence that optimizes the composite pseudo-log-likelihood of the native sequence with label smoothing of 0.1.

Caliby was trained for 30,000 steps with 4 gradient accumulation steps and a batch size of 8, taking about 20 hours on a single NVIDIA H100 GPU with 80 GB of RAM. We kept an exponential moving average (EMA) of model parameters with a decay of 0.99 to use for evaluation. For Caliby-L, we increased the number of encoder and decoder layers to 5, the hidden dimension to 256, and trained for 100,000 steps, with all other settings unchanged.

#### SolubleCaliby

SolubleCaliby was trained identically to Caliby, but with transmembrane proteins excluded from the training set, following SolubleMPNN^21^. We excluded all transmembrane PDB codes from SolubleMPNN’s exclusion list provided on the ProteinMPNN GitHub at https://github.com/dauparas/ProteinMPNN/blob/main/soluble_model_weights/excluded_PDBs.csv. We additionally excluded any transmembrane PDB codes annotated in the Protein Data Bank of Transmembrane Proteins at https://pdbtm.unitmp.org/downloads^43^.

#### CalibyPacker

For sidechain packing, we trained CalibyPacker using the same architecture as Caliby with the per-residue diffusion approach introduced in FAMPNN^7^. CalibyPacker uses 10 MPNN encoder and decoder layers at hidden dimension 128. The trunk is run once to produce per-residue node embeddings, which condition a lightweight per-token diffusion MLP that denoises sidechain heavy-atom coordinates in each residue’s local backbone frame. The diffusion head follows the EDM parametrization^44^ with self-conditioning implemented as described in FAMPNN. We trained for 100,000 steps with post-hoc EMA selection based on the validation set. We release checkpoints at backbone noise levels of 0.0Å and 0.1Å. Training and evaluation details closely follow FAMPNN, where performance on CASP13, 14, and 15 targets held out at 40% sequence similarity is reported in Supplementary Fig. 1.

### Ensemble design details

Given an ensemble of K structures, we first compute the neural-network-derived Potts model independently for each structure indexed by K:

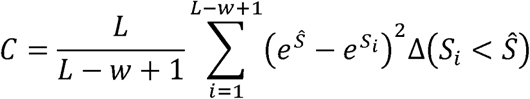

where Z_k_ is the normalizing factor known as the partition function. In practice, we also transform each Potts model to zero-sum gauge so that:

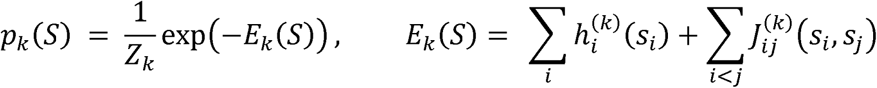

where A is the alphabet of amino acids. Then, given K input structures with Potts energies E_k_, we define the ensemble energy as the average:

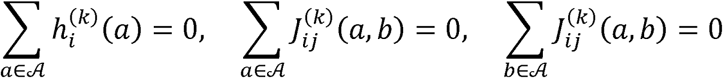

When designing on ensembles of K structures, we include the original structure as the first backbone and use partial diffusion structures for the remaining K - 1 structures.

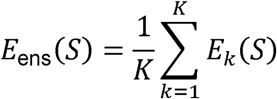

Because the ensemble-conditioned energy is defined as the average of the per-structure Potts energies, scoring a sequence under the ensemble is equivalent to scoring under a single Potts model with sitewise and pairwise energy terms averaged over the ensemble (Supplementary Note 1). As a result, in practice, Caliby precomputes a single ensemble-conditioned Potts model by averaging Potts models across structures, then samples sequences from it directly.

### Datasets

#### Training datasets

Our dataset curation closely follows the Boltz-1 procedure^45^. For our training set, we used all PDB structures released before 2021-09-30 with a resolution less than 9Å. We developed methodology using the validation set curated by Boltz-1, which uses MMseqs2^46^ to cluster sequences at a 40% sequence identity threshold to ensure that no sequences in the validation set fall into any train clusters. Then, the validation set is filtered for structures with resolution below 4.5Å and released between 2021-09-30 and 2023-01-13. The test set is constructed similarly to the validation set, but only using structures released after 2023-01-13.

For data preprocessing and data loading, we adopted the AtomWorks framework^47^ to parse the first bioassembly of each PDB and clean RCSB structures. During training, we sampled protein chains weighted by the inverse of the size of the cluster that the chain falls into and applied random contiguous cropping to the chain if the chain is longer than 1024 residues. We found that despite being trained on single chains, Caliby generalizes well to sequence design on protein complexes. From our preliminary experiments, we did not find evidence that training on multiple chains measurably improved *in* silico performance despite testing various modes of spatial cropping along interfaces, so we chose to simplify training and leave explicit interface training for future work.

#### Native backbone test set

To select native backbones for evaluation, we used the first bioassembly for each structure in the Boltz-1 test set and extracted all protein monomers between lengths 32 and 512, yielding 194 protein chains. We excluded 5 chains due to issues with consistent parsing between different sequence design methods, resulting in 189 chains for evaluation.

#### Backbone selection for experimental validation

To select candidate backbones for experimental characterization, we filtered the test set for monomeric, single domain apo structures. We excluded structures with chain breaks of 10 residues or longer, yielding 32 backbones. Among these, 5 had smaller chain breaks, which we infilled with Protpardelle-1c motif scaffolding.

### *In silico* evaluation and analysis

#### AlphaFold2 prediction

Unless explicitly stated, for all AlphaFold2 evaluations, we used AlphaFold2 in single-sequence mode with 3 recycles, taking the best by pLDDT out of 5 models, as implemented in ColabDesign^48^.

#### Designability evaluations

We ran ProteinMPNN sequence design with the 0.2Å checkpoint (vanilla_model_weights/v_48_020.pt) and all other sampling parameters at their defaults. To run ChromaDesign, we used the publicly available weights and all sampling parameters at their defaults (t=0.5). To run ESM-IF1^2^, we used the publicly available model weights from the official GitHub repository at https://github.com/facebookresearch/esm/tree/main.

To evaluate on RFdiffusion *de novo* backbones, we generated 100 RFdiffusion-generated backbones for each length in {100, 200, 300, 400, 500}, sampled one sequence per backbone from each sequence design model, then predicted with single-sequence AlphaFold2.

For evaluating Protpardelle-1c designs, we used the publicly available samples generated for the BindCraft benchmark at https://zenodo.org/records/17096818. For evaluating on BindCraft designs, we took a random subsample of 100 passing BindCraft trajectories per target and re-designed non-interface residues using either SolubleCaliby or SolubleMPNN with the same settings as in BindCraft. Following Lu et al.^13^, we predicted designs using single-sequence AlphaFold2 and used the template of the target, removed the binder template, removed interchain template features, did not use initial guess, and removed target template sequence features to allow for target backbone flexibility.

#### Sequence covariation analysis

To analyze whether ensemble conditioning reduces sequence covariation unrelated to structural constraints, we filtered our native backbone test set to remove structures with gaps and missing residues, yielding 111 structures. We then designed 1 sequences per structure with SolubleCaliby-fb and SolubleCaliby-e32. For computing categorical Jacobians, we used the ESM-2 3B parameter model^27^. Following Zhang et al.^26^, for each position, we mutated each position to each possible amino acid and recorded the change in output logits, yielding a [L, 20, L, 20] matrix, then mean-centered and symmetrized the result. Coupling scores were obtained by applying the Frobenius norm followed by average product correction. For interpolating between AlphaFold2 with full MSA and AlphaFold2 with only query sequence, we first generated MSAs for each designed sequence and predicted the structure using ColabFold^49^. We then randomly masked proportions of columns in the MSA in increments of 5% and ran structure prediction with the partially masked MSA. For each masking proportion, we randomly masked with 3 different random seeds and averaged results across seeds.

For direct-coupling analysis experiments, we filtered the test set to remove sequences whose MSA had fewer than 512 sequences in addition to removing structures with gaps and missing residues, yielding 96 structures. To subsample natural MSAs to 512 sequences, we used the greedy diversity maximization strategy implemented by Akiyama et al. for MSA Pairformer^25^. We inferred contacts using Markov random field pseudo-likelihood maximization by running CCMpred (https://github.com/soedinglab/CCMpred)^50^. We defined a residue pair as a contact if their Cβ atoms (Cα for glycine) were within 8Å, and a long-range contact as a contact at least 24 residues apart in sequence space. For a protein of length L, long range contact precision at L (P@L) is defined as the fraction of the top L predicted contacts that are true long-range contacts. For the MSA Pairformer experiments, we used the same set of 96 structures, contact definitions, and synthetic MSAs, but provided the native sequence as the query to infer contacts for it.

#### Native sequence bias for interpolation paths

To generate interpolation paths between full Caliby redesign and native-like sequences, we introduced a soft bias toward the native amino acid sequence during Potts sampling. For each protein, after computing the fixed-backbone-conditioned or ensemble-conditioned Potts model with sitewise fields h_i_(a) and pairwise couplings J_ij_(a,b), we biased the field of the native amino acid at every position:

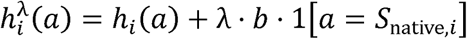

where λ controls the bias strength and b is a protein-specific scale. This modifies only the sitewise fields, leaving pairwise couplings unchanged. At λ = 0, sampling is equivalent to unbiased design with Caliby, while larger λ results in greater retention of native residues.

Because Potts energies are unnormalized across proteins, we chose b to make λ comparable across proteins. Specifically, we first evaluated the local conditional energies of the native sequence under the unmodified Potts model:

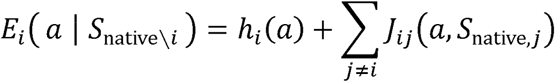

This scores how energetically favorable an amino acid a is at position i given the native amino acid sequence at all other positions. Then, we defined b as:

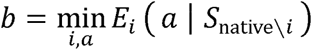

In other words, b is the most favorable local conditional energy across all positions and amino acid types. This scales the bias so that at λ = 1, each position receives a bias towards the native residue type comparable to the strongest single-residue preference in the Potts model.

For interpolation experiments, we swept λ from 0.0 to 1.0 in increments of 0.02, sampling 8 sequences per backbone with either SolubleCaliby or SolubleCaliby-e32 while excluding cysteines from sampling. We performed experiments on the same set of 32 monomeric, single domain apo structures used for selecting experimental candidates, excluding 5 structures with missing residues (n=27).

#### Local energetic frustration

To quantify local energetic frustration, we used Frustratometer^11,12^ to compute frustration indices under the AWSEM energy function^28^. In practice, we used the Python implementation of Frustratometer at https://github.com/HanaJaafari/Frustratometer. In Fig. 4, we report the single-residue mutational frustration index, which evaluates how favorable the native amino acid at each position is relative to a set of decoy substitutions, with the structure and all other residues held fixed. In Extended Data Fig. 6, we report the configurational frustration index, which is computed over pairwise contacts and randomizes both residue identities and their spatial arrangement. Both frustration indices are defined as the Z-score of the native energy relative to the decoy distribution. Following Ferreiro et al.^11^, we define residues or contacts with frustration index below -1 as highly frustrated and above 0.78 as minimally frustrated. We define average frustration as the negated frustration index averaged across residues or contacts, such that higher values correspond to greater frustration.

#### Lanthanide binder design

Lanthanide-binding backbones were obtained from a separate in-house design campaign. Briefly, Protpardelle-1c was used to scaffold the four EF-hand loops of the lanmodulin from *Hansschlegelia quercus* (*Hans*-LanM; PDB: 8DQ2)^32^. ProteinMPNN was used to design sequences, with the two strongest-binding loops, EF2 and EF3, held fixed to the native loop sequence. Designs were co-folded with two lanthanide ions and filtered by Boltz-1 confidence^45^, yielding 34 backbones of length 100 that adopt the *Hans*-LanM fold, with TM-scores around 0.75 to 0.85 to the native.

For design with Caliby, we designed 8 sequences per backbone with SolubleCaliby-e32, excluding cysteines and holding the EF2 and EF3 loop positions fixed. Each designed sequence was co-folded with 2 Sm^3+^ ions using Boltz-2^51^, and we retained the best sequence by confidence score for each backbone. We then ranked backbones by the best composite score achieved by either SolubleCaliby-e32 or ProteinMPNN. Among the top 10 backbones, we chose an arbitrary set of 5 SolubleCaliby-e32 designs for experimental characterization.

#### Homo-oligomeric membrane protein design

We selected homo-oligomeric membrane protein assemblies from QSProteome^34^, a community-led repository of predicted quaternary structures, with predicted interface TM-score ≥ 0.8, model quality score ≥ 0.8, and total assembly size ≤ 1800 residues. From these criteria, we chose 8 membrane targets spanning a range of 3-to 10-fold symmetry (Supplementary Table 2). We ran SolubleCaliby-e32 on the complete assembly, excluding cysteines and tying positions symmetrically across chains during sampling, following the implementation in ChromaDesign. Briefly, symmetry is enforced by summing the Potts fields and couplings for all symmetry-equivalent positions into a single representative position, sampling over the reduced position set, and then copying each sampled identity back to its tied group.

Designs were evaluated against the full assembly by self-consistency using single-sequence AlphaFold2 in homo-oligomer mode and AlphaFold-Multimer^35^. We screened targets by first designing 8 sequences per target, advancing any target with a design with scTM of at least 0.7 under either predictor, yielding 4 of 8 targets. Then, we designed an additional 112 designs for these 4 targets. Among all 120 designs for each of these targets, we filtered for mean pLDDT ≥ 75 under both predictors. Among this set of filtered designs, we selected the design maximizing the minimum pLDDT across the two for experimental characterization.

### Experimental characterization of designs

#### Protein expression and purification

Genes encoding the designed protein sequences were synthesized and cloned into pET-24a(+) *Escherichia coli* plasmid expression vectors containing a C-terminal 6xHis-tag (Genscript). Plasmids were transformed into chemically competent BL21(DE3) *E.□coli* cells (Zymo Research), which were cultured in 2xYT medium at 37°C until the optical density (OD) reached 0.6–0.8. Protein expression was induced with 1□mM IPTG, and cultures were incubated overnight at 16□°C. Cells were harvested by centrifugation, resuspended in 50□mM Tris-HCl, 150□mM NaCl (pH 8.0), and frozen at −80□°C until purification. Frozen cell pellets were thawed and lysed by sonication. The spun down supernatants of the lysates were filtered and equilibrated with 20 mM sodium phosphate (pH 7.4), 300 mM NaCl, 10 mM imidazole before incubation with Ni-NTA resin (Thermo Scientific, 88223) for 1 h at room temperature with batch binding. The resin was washed twice using 20 mM sodium phosphate (pH 7.4), 300 mM NaCl, 25 mM imidazole, and bound proteins were eluted in three fractions using 20 mM sodium phosphate (pH 7.4), 300 mM NaCl, 250 mM imidazole. Eluted proteins were buffer-exchanged into 50 mM Tris-HCl, 150□mM NaCl (pH 8.0), concentrated, and further purified by size-exclusion chromatography using either a Superdex 75 10/300GL or Superdex 200 10/300GL (Cytiva). Purified protein samples were analyzed by SDS– PAGE (BioRad). Protein concentrations were determined by measuring absorbance at 280□nm with a Nanodrop spectrophotometer (Thermo Fisher Scientific) using the predicted extinction coefficient and molecular weight.

#### Circular dichroism and thermal denaturation

CD spectra were measured on a JASCO CD spectrophotometer in a 1-mm-pathlength cuvette (Hellma). Protein samples were at ∼0.2□mg□ml−1 in the 50□mM TrisHCl 150mM NaCl, (pH 8.0) buffer. Melting temperature ranged from 20 to 95□°C and the absorption signal was monitored at a wavelength between 211 nm and 221□nm in 1□°C increments per minute, with 10□s of equilibration time and 1□s of digital integration time. Wavelength scans (200–260□nm) were collected at 20 and 95□°C and again at 2□°C after fast refolding.

#### Purification of Lanthanide Binders

Genes encoding the designed protein sequences were synthesized and cloned into pET-24a(+) Escherichia coli plasmid expression vectors (Genscript), wherein the protein sequence contained a C-terminal Strep-tag. Plasmids were transformed into BL21(DE3) competent E. coli cells. These cells were grown in 1L of 2XYT media at 37°C until an OD of 0.6-0.8 was reached. Cultures were then centrifuged (4,000g, 4°C, 30 minutes), and the supernatant was discarded. The pellet was resuspended in Buffer AT (50mM Tris, 150 mM NaCl, pH 8.0) and stored at -20°C until lysis and purification.

To lyse the cells, PMSF in isopropyl alcohol was added to a final concentration of 1mM to the thawed resuspension. The cells were sonicated (20 seconds on, 40 seconds off, amplitude 20%, 9.5 minutes), then centrifuged (18,000g, 4°C, 1 hour). The supernatant was collected for purification via Strep-Tactin affinity columns followed by size-exclusion chromatography.

Purification was performed with Strep-Tactin. 4Flow. high-capacity resin, acquired from IBA LifeSciences. Before each use, columns were washed with at least 10 CV Milli-Q water, 10 CV 100mM NaOH, and 10 CV Milli-Q water. They were then equilibrated 2x with at least 10 CV Buffer AT. The cell lysate was added to the column and allowed to flow through. The column was then washed with 4 CV Buffer AT and 0.5 CV Buffer E. Protein was then eluted from the column 5x with 0.5 CV Buffer E, and these 5 fractions were collected for subsequent purification via SEC. To clean the Strep-Tactin columns, columns were washed 2x with at least 10 CV Milli-Q water, 2x with at least 10 CV 100mM NaOH, then 2x with at least 10 CV Milli-Q water again, after which they were stored in 20% ethanol.

The 5 elution fractions from the Strep-Tactin columns were combined and concentrated to 1mL using Amicon 3kDa MWCO Centrifugal Filter Units. The concentrated protein solutions were then purified via SEC (Superdex 75 10/300 GL), where Chelex-treated Buffer XO6 (20mM MES, 100mM KCl, pH 6.0) was used as a running buffer. All Chelex-treatment was done with 10g/L of Chelex-100 (Sigma-Aldrich, C7901) overnight.

#### Xylenol Orange Binding Assays

Metal-binding of the designed proteins was assessed in 96-well plates through a competition assay with xylenol orange (XO) dye as an indicator, which changes from yellow-orange to purple upon binding free metal^33,52^.

XO dye (Sigma-Aldrich, 52097) was first dissolved in Chelex-treated Milli-Q water to a stock concentration of 5mM and stored in -20°C. Samarium stock solution was obtained at a concentration of 10.014g/L (confirmed through ICP-MS) and stored at room temperature.

To prepare wells, all stock solutions were then diluted to their working concentrations in Chelex-treated Buffer XO6. Samarium stock was diluted down to 500uM, and metal samples ranging from 50 to 450 uM were prepared from this stock. XO dye stock was diluted by a factor of 450. Protein concentrations were determined using a Nanodrop at 280nm and were then diluted down to a working concentration of 10uM in Buffer XO6.

To each well, 45μL XO dye, 50μL protein of interest, and 5μL metal was added, so that the final wells contained 100μL total of 5uM XO, 5uM protein, and metal concentrations ranging from 0 to 25uM. For the standard curve, 50μL Buffer XO6 was used in place of protein. Solutions in each well were mixed by pipetting, and well absorbances were read at 574nm using a SpectraMax iD3 Multi-Mode Microplate Reader.

#### Mass photometry

Fractions corresponding to the target peak from SEC were pooled and concentrated for mass photometry analysis. Concentrated samples were diluted to 100 nM in 50 mM Tris-HCl, 150 mM NaCl (pH 8.0) and further diluted directly on a TwoMP MassGlass slide (Refeyn) to final concentrations of 10 nM using a 6-well cassette. Protein landing events were recorded using AcquireMP software (Refeyn) and analyzed with DiscoverMP software (Refeyn).

#### Cryo-EM sample preparation

The Caliby-designed Q8XAZ3 protein was purified and diluted to 6 mg/mL in 50 mM Tris-HCl, 150 mM NaCl (pH 8.0) supplemented with 0.02% amphipol A8-35 (Anatrace, A835). A 3.8 μL aliquot was applied to a glow-discharged (60 s; 0.38 mbar, 10 mA; PELCO easiGlow) UltrAuFoil R1.2/1.3 300-mesh gold TEM grid (Electron Microscopy Sciences, cat. no. Q350AR13A). Grids were plunge-frozen in liquid ethane using a Vitrobot Mark IV (Thermo Fisher Scientific) with a blot time of 4 s, blot force of 4, at 22□°C and 100% relative humidity.

#### Cryo-EM data collection, processing, and data analysis

Frozen grids were loaded into a 200 kV Glacios transmission electron microscope (Thermo Fisher Scientific) equipped with a Falcon 4i direct electron detector. Movies were collected at a nominal magnification of 150,000×, corresponding to a calibrated pixel size of 0.93 Å, with a total electron dose of 40.1 e^-^/Å^2^ and a defocus range of −0.9 to −2.7 μm. Automated data collection was performed using EPU (Thermo Fisher Scientific). A total of 728 movie stacks were collected, of which 707 were retained for motion correction, contrast transfer function (CTF) estimation, particle picking, 2D classification, *ab initio* reconstruction, heterogeneous refinement, homogeneous refinement, reference-based motion correction, and non-uniform refinement in cryoSPARC v4.7.1^53^. The final reconstruction was sharpened with EMReady2^54^. A schematic of the cryo-EM data processing workflow is shown in Extended Data Fig. 10. An AlphaFold3-predicted model was used as the initial model for atomic model building and was manually adjusted in Coot-1.1.10^55^, refined in ISOLDE^56^ implemented in ChimeraX v1.10.1^57^, and subjected to real-space refinement in Phenix-2.1-6048^58^. Cryo-EM data collection, refinement, and validation statistics are summarized in Supplementary Table 1.

## Data availability

The structural model generated in this study has been deposited in the Protein Data Bank (PDB) under the accession code of 38DW, and the corresponding cryo-EM map is available in the Electron Microscopy Data Bank (EMDB) under the accession code of EMD-78753.

## Code availability

Source code, sampling code, and training code for Caliby are available at https://github.com/ProteinDesignLab/caliby. Pre-trained model weights are hosted on HuggingFace at https://huggingface.co/ProteinDesignLab/caliby-weights. An interactive demo with free GPU access is available as a HuggingFace Space at https://huggingface.co/spaces/ProteinDesignLab/caliby, and a tutorial Google Colab notebook is provided in the GitHub repository.

## Acknowledgements

We thank Alex Chu and Sergey Ovchinnikov for discussion and Martin Pacesa for providing the raw BindCraft trajectories used for evaluation. Computation was performed on the Sherlock and Marlowe clusters at Stanford University. Cryo-EM data were collected at the Stanford-SLAC Cryo-EM Center. R.W.S. acknowledges funding support from the NSF Graduate Research Fellowship (DGE-2146755). T.L. is supported by the Stanford Graduate Fellowship. P.K. acknowledges the computational resources provided at the LUMI supercomputer owned by the EuroHPC Joint Undertaking, hosted by CSC (Finland) with access granted through IT4I National Supercomputing Center, Czech Republic and the e-INFRA CZ (No. 90254) project. The research stay of P.K. at Stanford was funded by the European projects CLARA (No. 101136607), ERC project FRONTIER (No. 101097822), COST Action CA21162 COZYME as well as by G-Research, Loschmidt Laboratories, Czech Technical University in Prague and Masaryk University. Additional support to P.-S.H came from Merck Research Laboratories (MRL) Scientific Engagement and Emerging Discovery Science (SEEDS) Program, Stanford Medicine Catalyst, and NIH (R01GM147893).

## Author contributions

Conceptualization: R.W.S. and P.-S.H.; methodology: R.W.S., T.L. and P.K.; software: R.W.S. and J.Y.; investigation: R.W.S., L.Z., G.E.N., J.L., T.L., I.M. and S.B.; formal analysis: R.W.S., L.Z. and G.E.N.; data curation: R.W.S.; writing—original draft: R.W.S., L.Z., G.E.N., J.L., T.L. and P.-S.H.; writing—review and editing: R.W.S., L.Z., G.E.N., J.L., T.L., I.M., S.B., J.Y., P.K. and P.-S.H.; visualization: R.W.S., L.Z., G.E.N. and J.Y.; supervision: P.-S.H.; funding acquisition: P.-S.H.

## Competing interests

The authors declare no competing interests.

## Additional information

Correspondence and requests for materials should be addressed to Po-Ssu Huang.

## Extended Data Figures

**Extended Data Figure 1.**
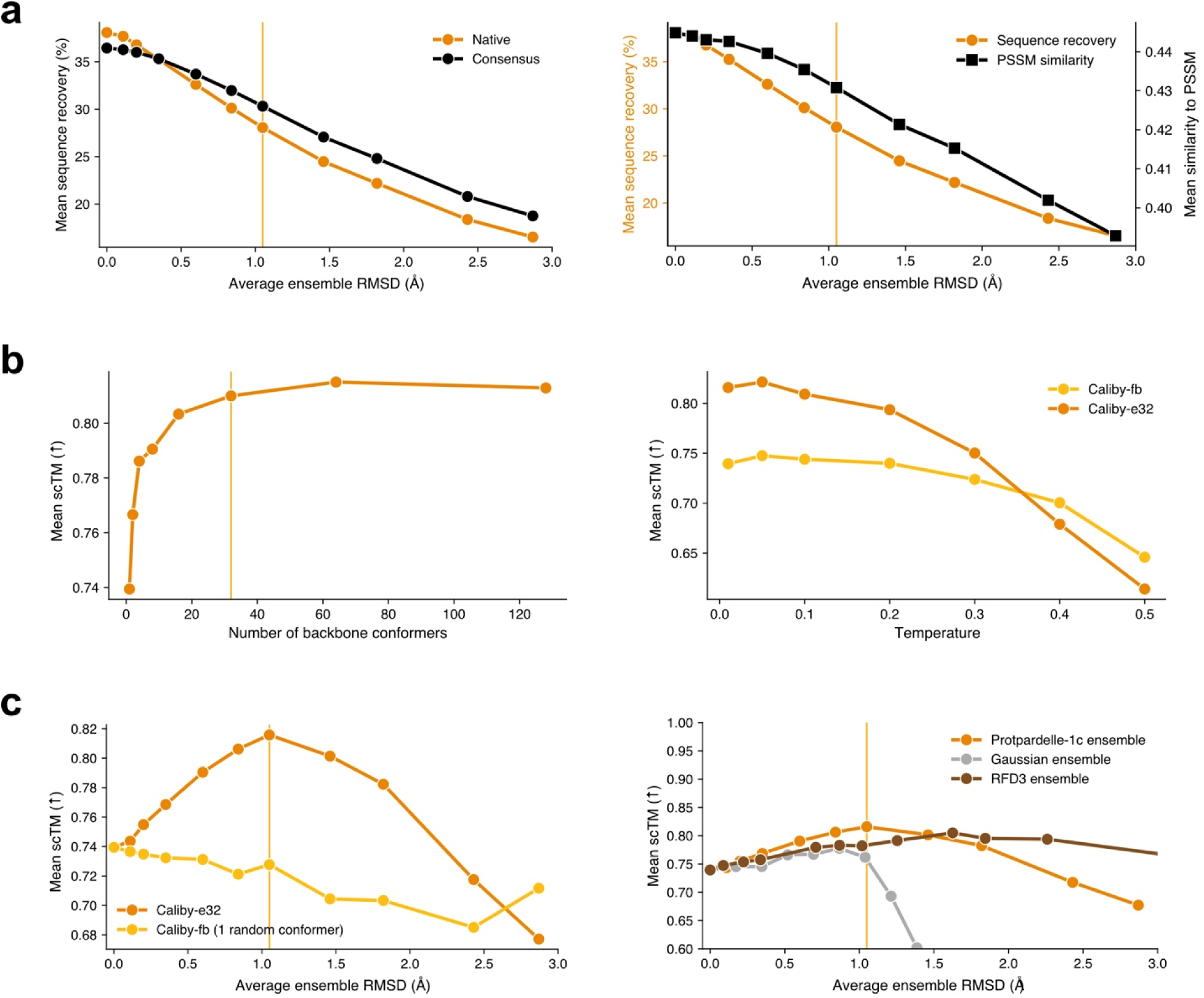
Ensemble conditioning ablations. **(a)** Native sequence recovery compared with consensus (left) and PSSM similarity (right) as a function of ensemble structural diversity. Consensus recovery and PSSM similarity decline more slowly than native sequence recovery. The orange vertical line denotes 150 rewind steps, used as the default. **(b)** Mean scTM as a function of ensemble size (left) and sampling temperature for Caliby-fb and Caliby-e32 (right). The orange vertical line denotes 32 conformers, used as the default. **(c)** Left, mean scTM for Caliby-e32 versus Caliby-fb run on a single random conformer from the synthetic ensemble. Because partial diffusion may improve designability by refining imperfect backbones, this ablation confirms that the improvement from ensemble conditioning is not attributable to backbone refinement alone. Right, mean scTM for ensembles generated by Protpardelle-1c, RFdiffusion3, and Gaussian noise as a function of ensemble structural diversity.

**Extended Data Figure 2.**
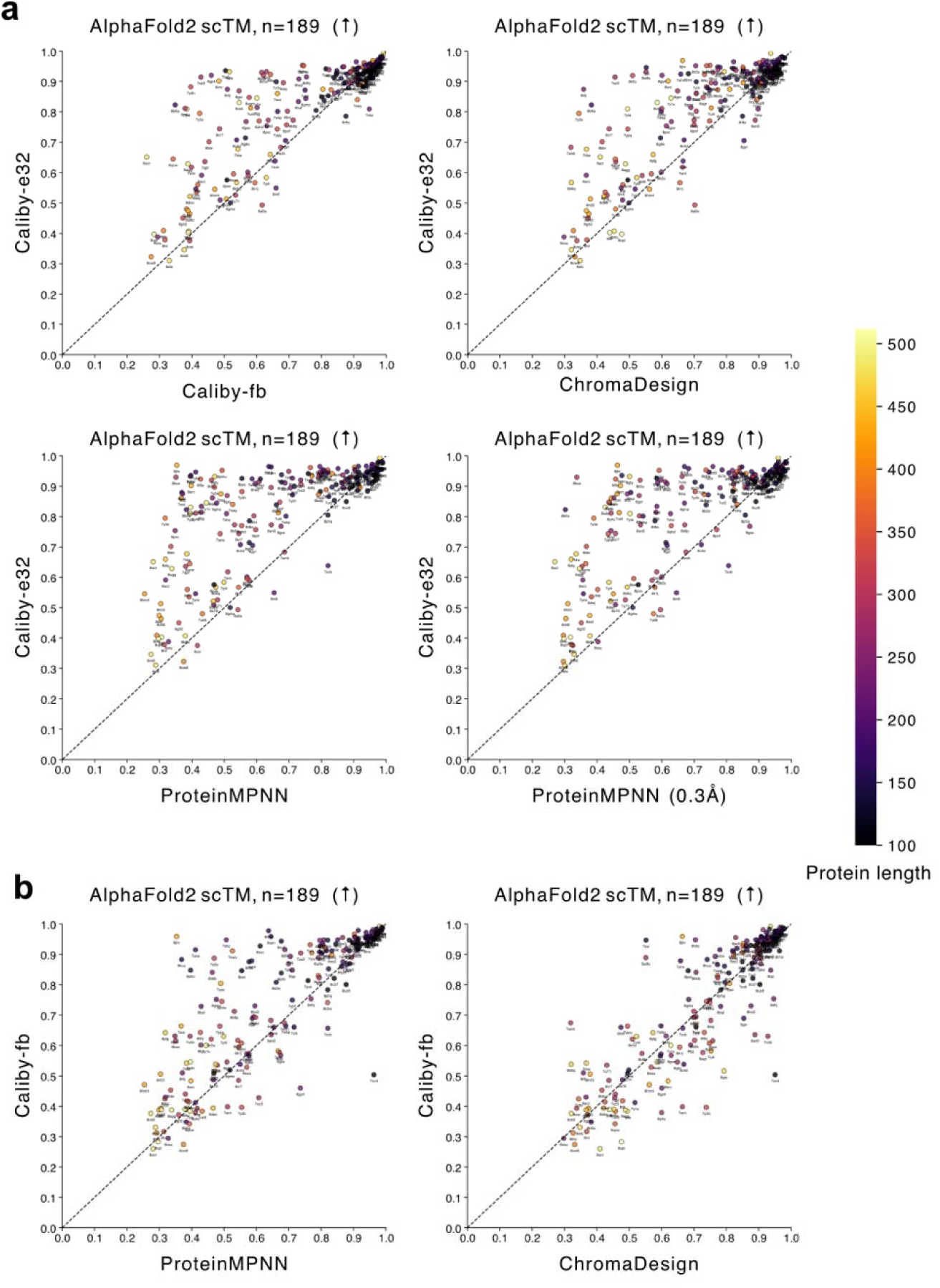
Native backbone test set scTM comparisons. Best of 8 designs shown per backbone, colored by protein length. **(a)** Single-sequence AlphaFold2 scTM for Caliby-e32 vs. Caliby-fb, ChromaDesign, ProteinMPNN, ProteinMPNN using the 0.3Å checkpoint (n = 189). **(b)** Single-sequence AlphaFold2 scTM for Caliby-fb vs. ProteinMPNN and ChromaDesign (n = 189).

**Extended Data Figure 3.**
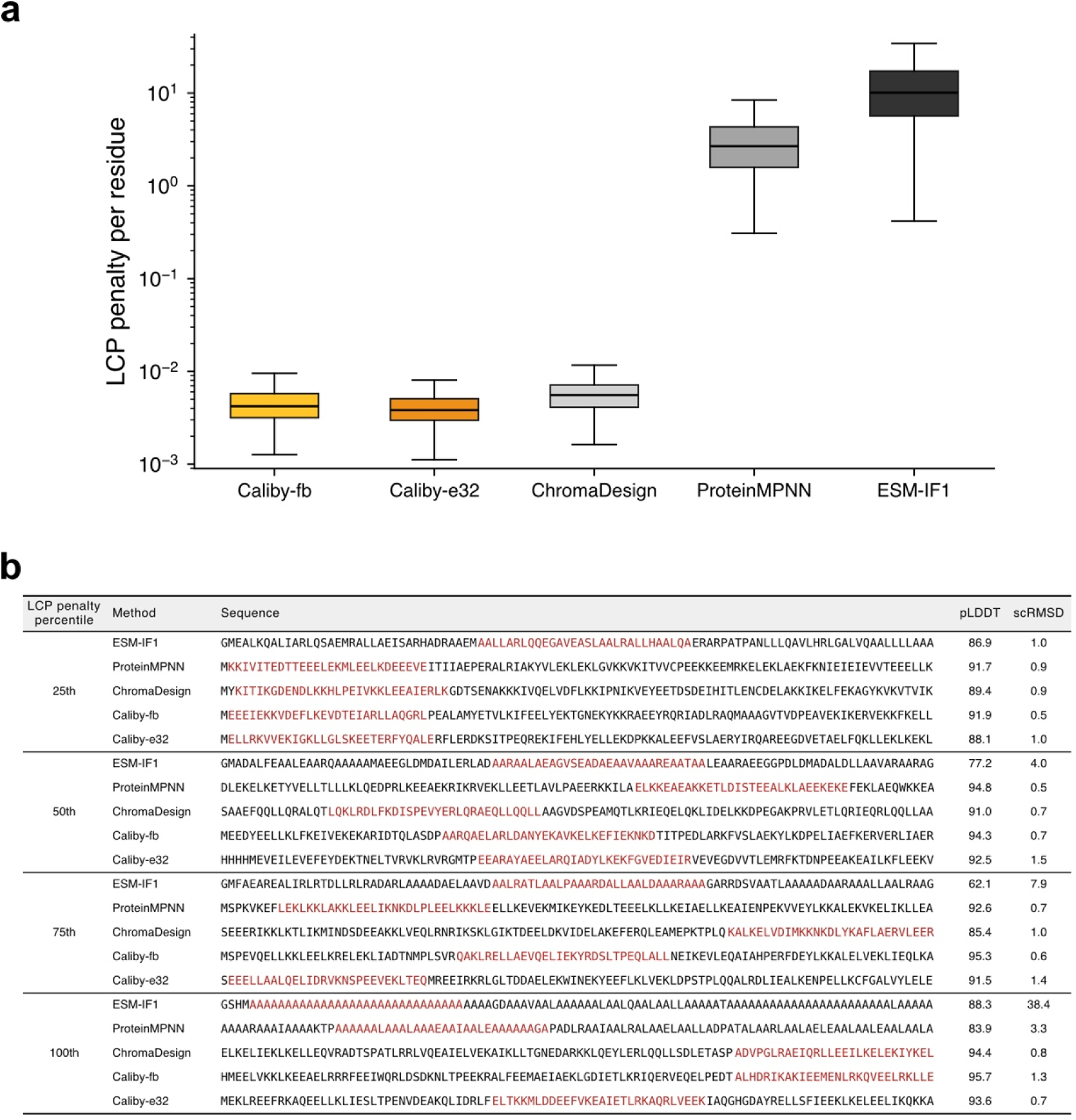
Sequence complexity across design methods. **(a)** Local composition perplexity (LCP) penalty per residue for sequences designed by each method on RFdiffusion backbones from Fig. 2b (Methods). Higher values indicate lower sequence complexity. ProteinMPNN and ESM-IF1 design sequences with higher LCP penalties, reflecting a tendency towards repetitive, low-complexity sequences. Caliby and ChromaDesign maintain low penalties due to an LCP restraint during sampling. **(b)** Example sequences at the 25th, 50th, 75th, and 100th percentiles of LCP penalty for RFdiffusion backbones of length 100, with the highest penalty 30-residue window highlighted in red. The 100th percentile ProteinMPNN sequence is nearly all-alanine yet achieves a pLDDT of 83.9.

**Extended Data Figure 4.**
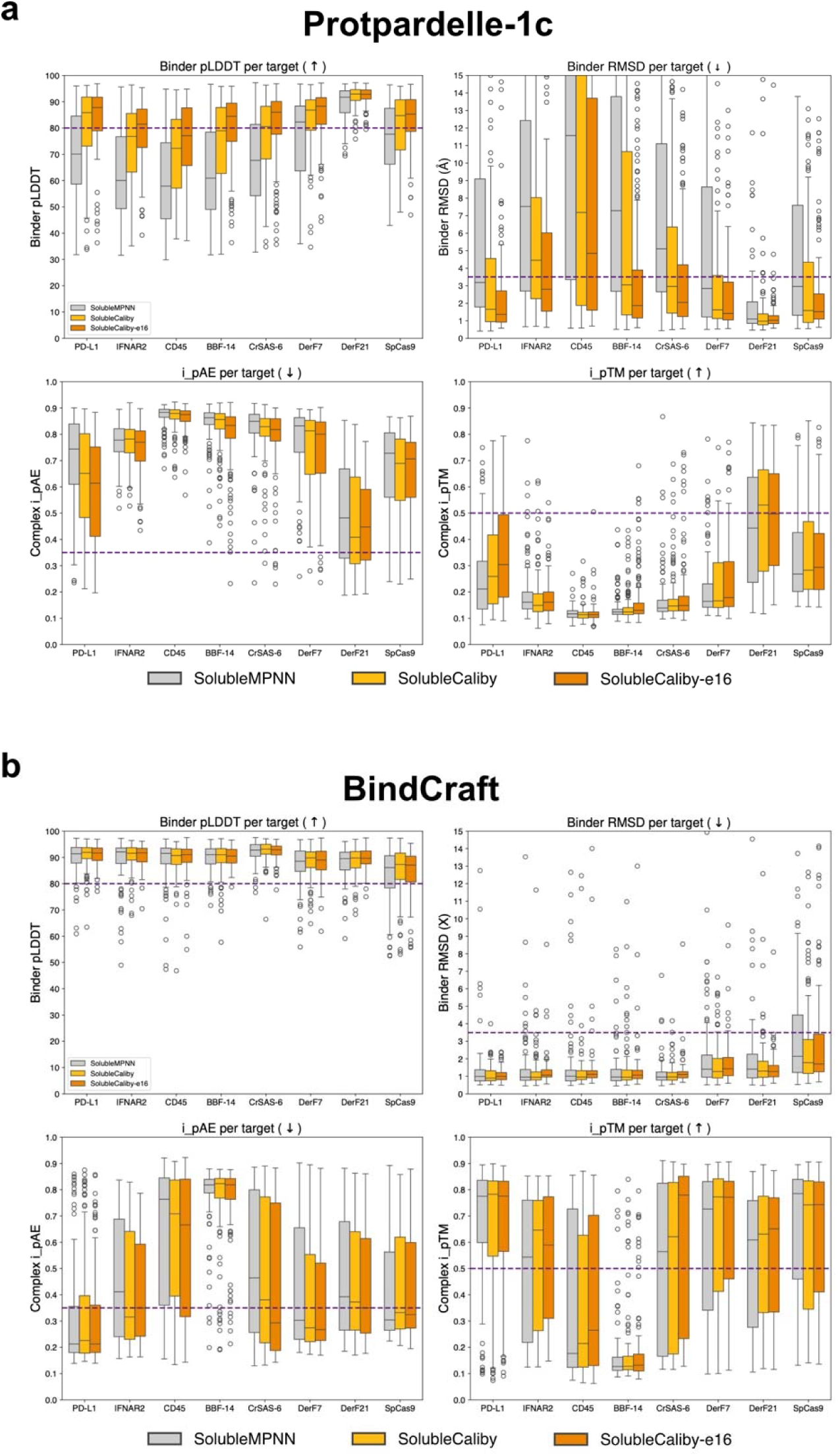
BindCraft binder design benchmark. **(a)** SolubleMPNN vs. SolubleCaliby vs. ensemble-conditioned SolubleCaliby used for designing non-interface residues on binders designed by Protpardelle-1c for various BindCraft targets. Success thresholds are denoted as purple dashed lines (i_pTM > 0.5, i_pAE < 0.35, binder pLDDT > 0.8, binder RMSD < 3.5Å). **(b)** Same as (a) but with BindCraft-generated binders instead of Protpardelle-1c-generated binders.

**Extended Data Figure 5.**
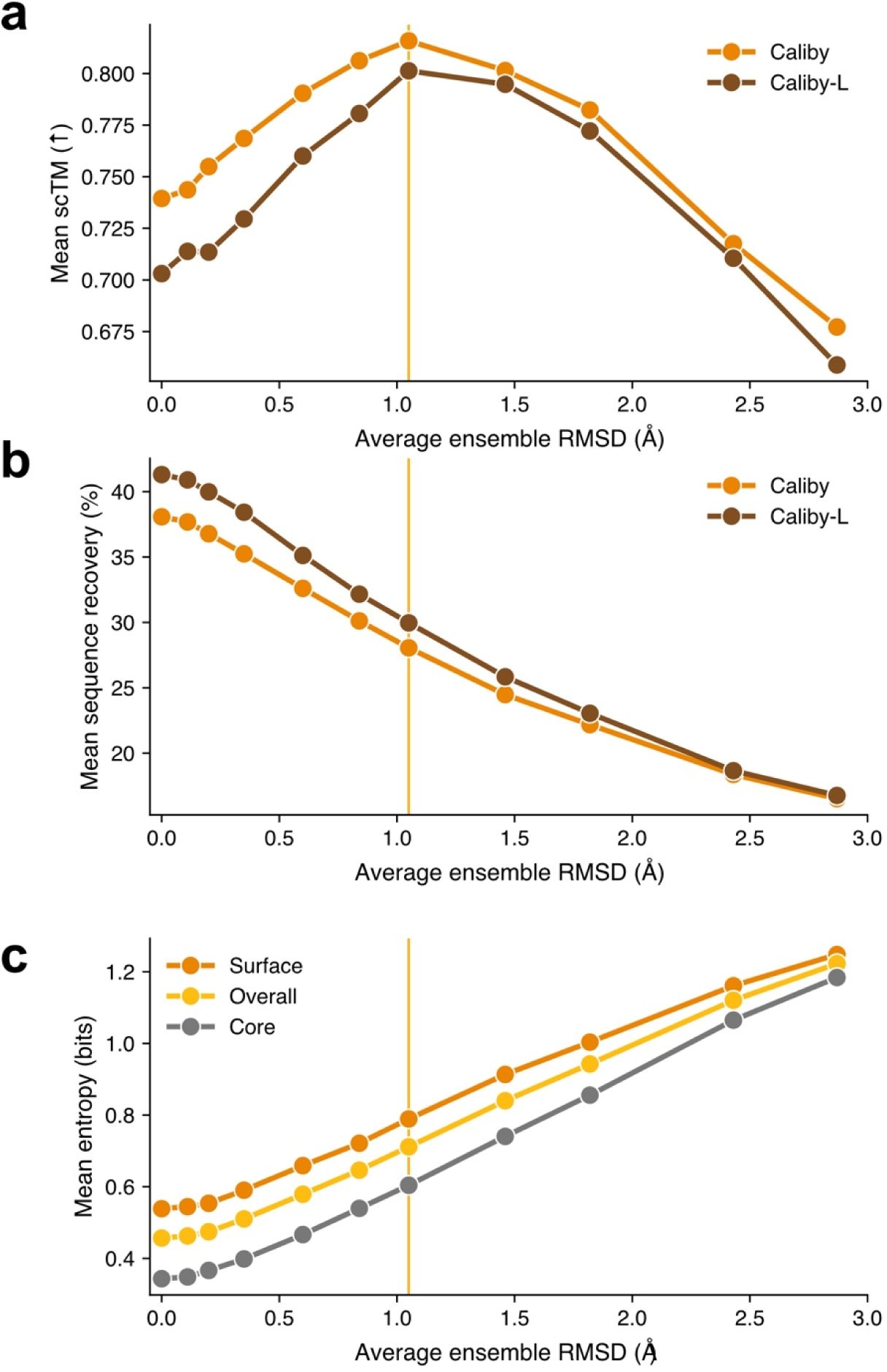
Effects of ensemble structural diversity on designed sequences. **(a)** Mean single-sequence AlphaFold2 scTM as a function of ensemble structural diversity for Caliby and Caliby-L, where Caliby-L is a larger model with 5 MPNN layers instead of 3 and a hidden dimension of 256 instead of 128. 0Å denotes fixed-backbone design, and the orange vertical line denotes 150 Protpardelle-1c rewind steps, used as the default. **(b)** Mean native sequence recovery over the same backbones. **(c)** Mean positional entropy of sequences designed by Caliby at surface, core, and all positions.

**Extended Data Figure 6.**
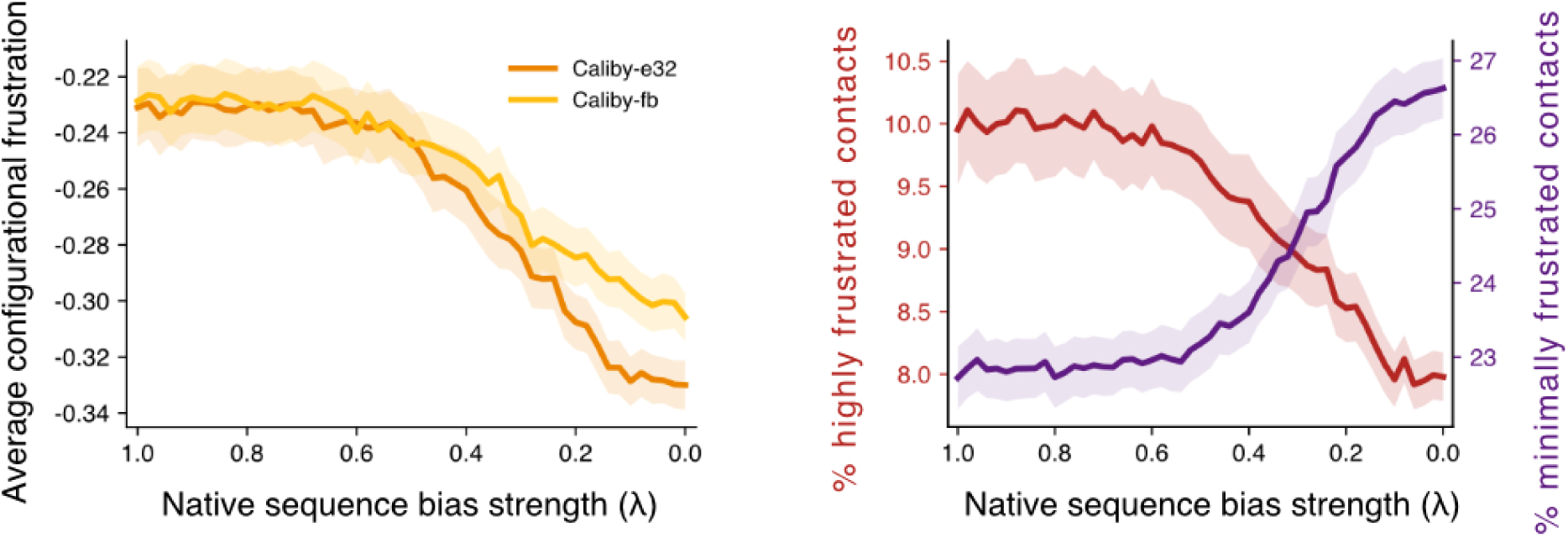
Configurational frustration along the native sequence interpolation path. **(a)** Average configurational frustration, defined as the negated configurational frustration index from Frustratometer, for Caliby-fb and Caliby-e32 as a function of native sequence bias strength λ. (b) Percentage of highly frustrated and minimally frustrated contacts for Caliby-e32. Shaded regions show ± 1 SEM.

**Extended Data Figure 7.**
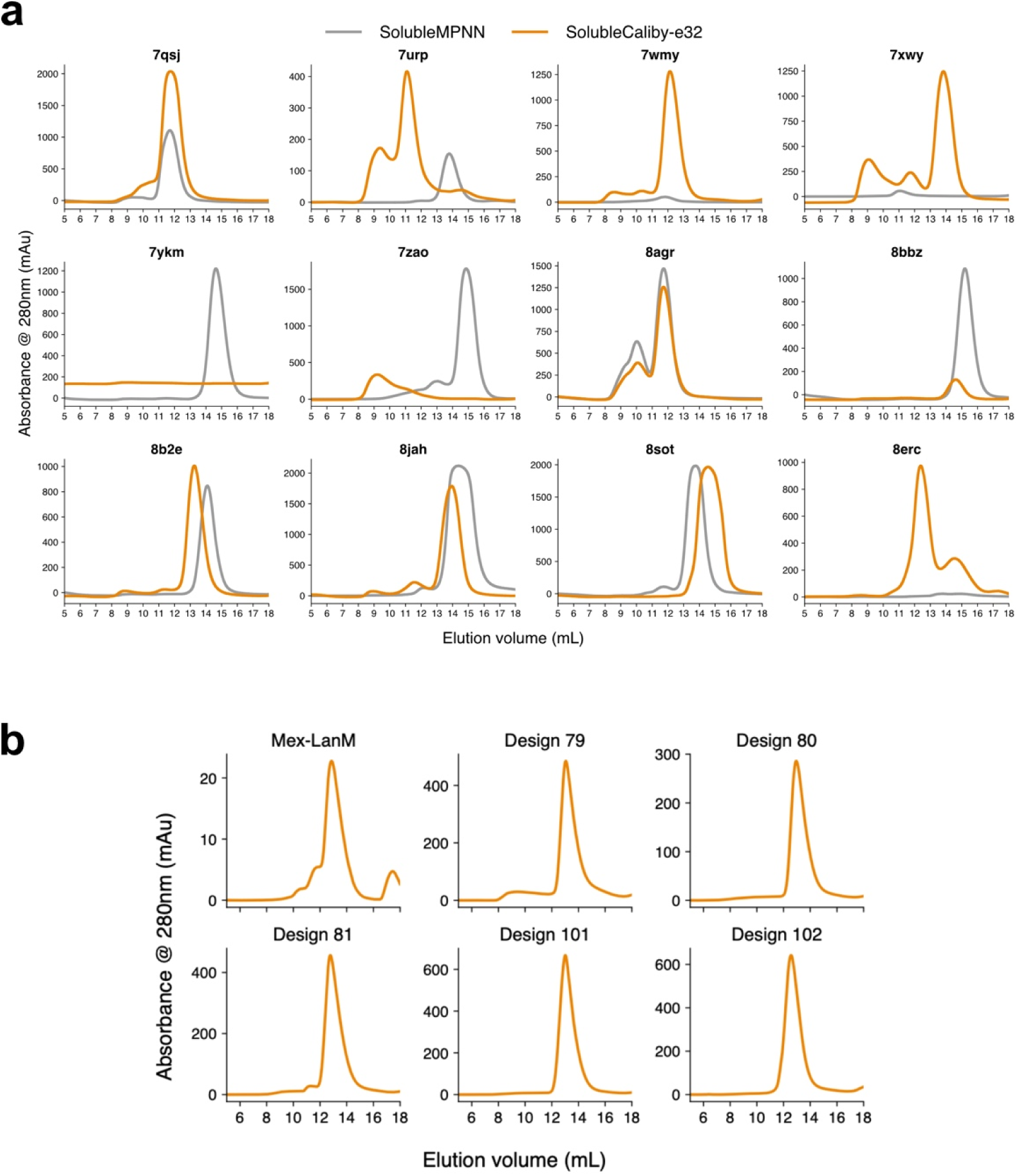
Size exclusion chromatography of native protein redesigns and lanthanide binders. **(a)** SEC traces (absorbance at 280 nm) for SolubleMPNN (gray) and SolubleCaliby-e32 (orange) designs across all 12 target native backbones, labeled by PDB code. The SolubleCaliby design for 7YKM contains no Trp or Tyr residues and has negligible predicted absorbance at 280 nm, so its elution is not detectable in the A280 SEC trace. The SolubleMPNN design for 8ERC was recovered predominantly in the insoluble fraction. **(b)** SEC traces for the native lanmodulin Mex-LanM and the 5 SolubleCaliby-e32 lanthanide-binding designs characterized in Fig. 5c.

**Extended Data Figure 8.**
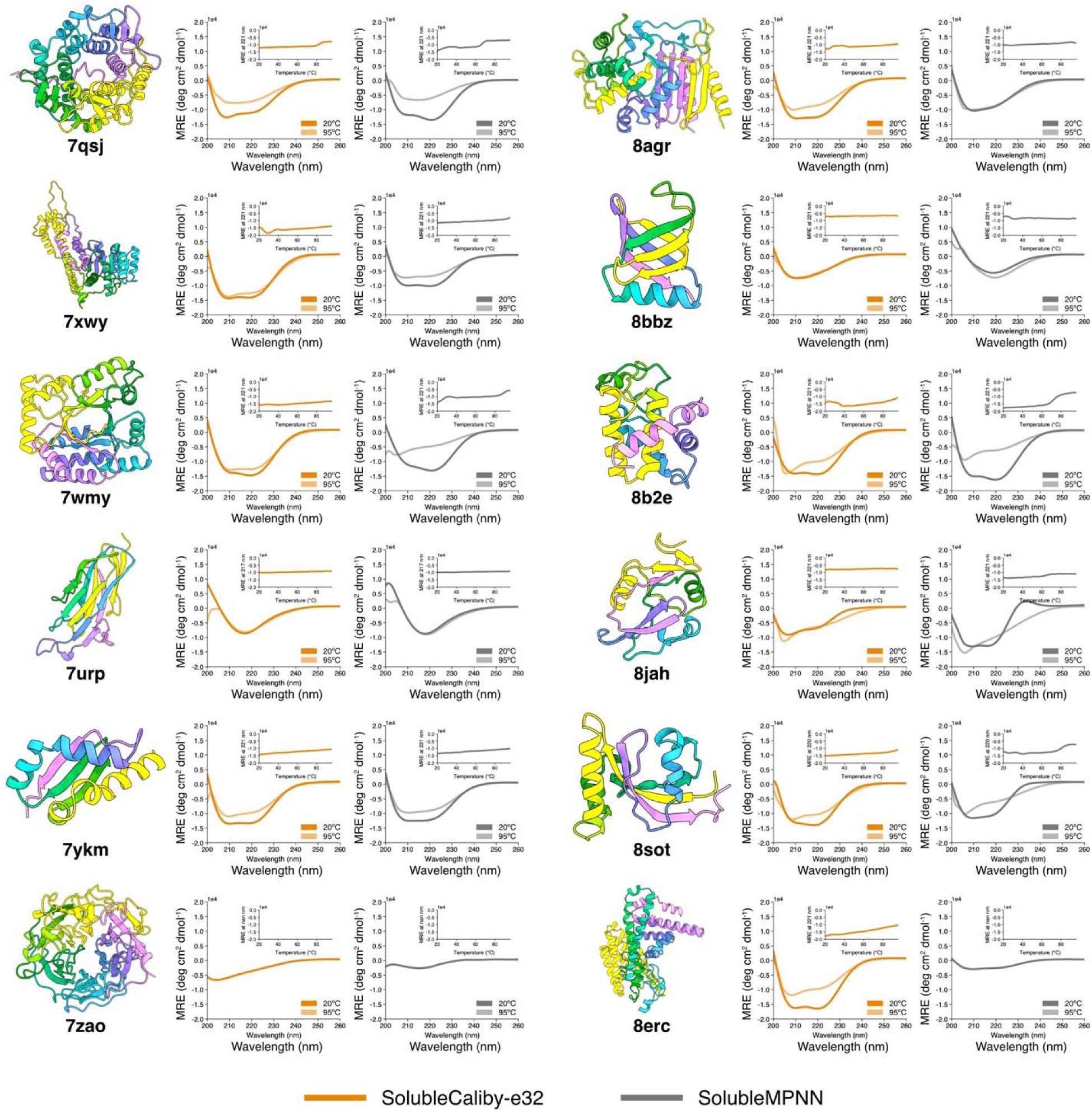
Circular dichroism spectra and thermal melts for native protein redesigns. Native backbone structures (rainbow) and CD wavelength scans (200-260 nm) at 20°C (dark) and 95°C (light) with thermal denaturation curves (insets) for SolubleCaliby-e32 (orange) and SolubleMPNN (gray) designs across all 12 target native backbones, labeled by PDB code.

**Extended Data Figure 9.**
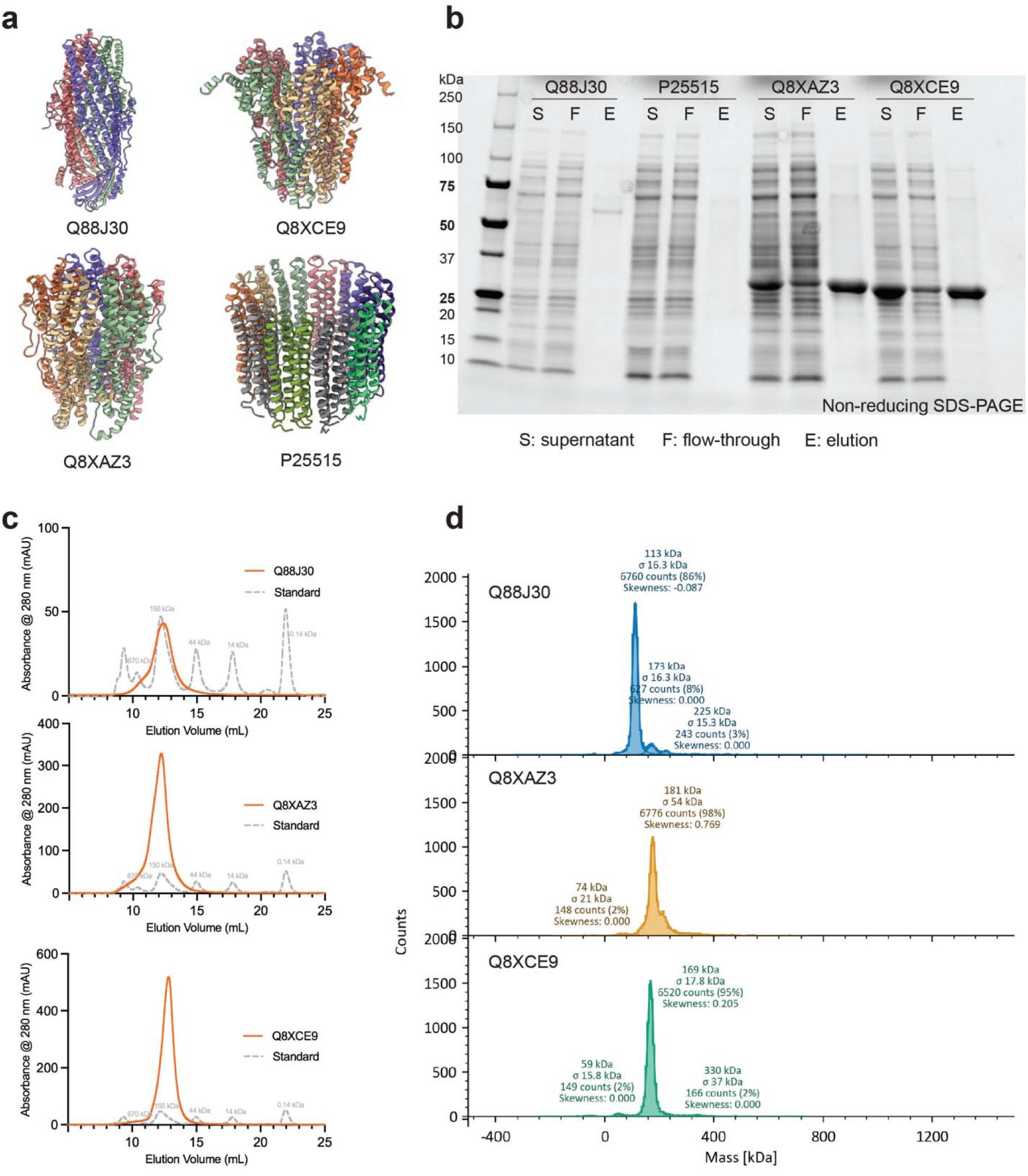
Purification and biophysical characterization of four Caliby sequence-designed homo-oligomeric protein assemblies. **(a)** AlphaFold3 predictions of the designed protein structures. **(b)** Ni-NTA affinity purification analyzed by SDS-PAGE, showing elution fractions containing proteins at the expected molecular weights. **(c)** Size-exclusion chromatography traces for each Caliby-designed construct on a Superdex 200 Increase 10/300 GL column (Cytiva). **(d)** Mass photometry (TwoMP, Refeyn) measurements of the SEC-purified fractions at 10 nM, showing the estimated molecular weight distributions of each designed protein assembly.

**Extended Data Figure 10.**
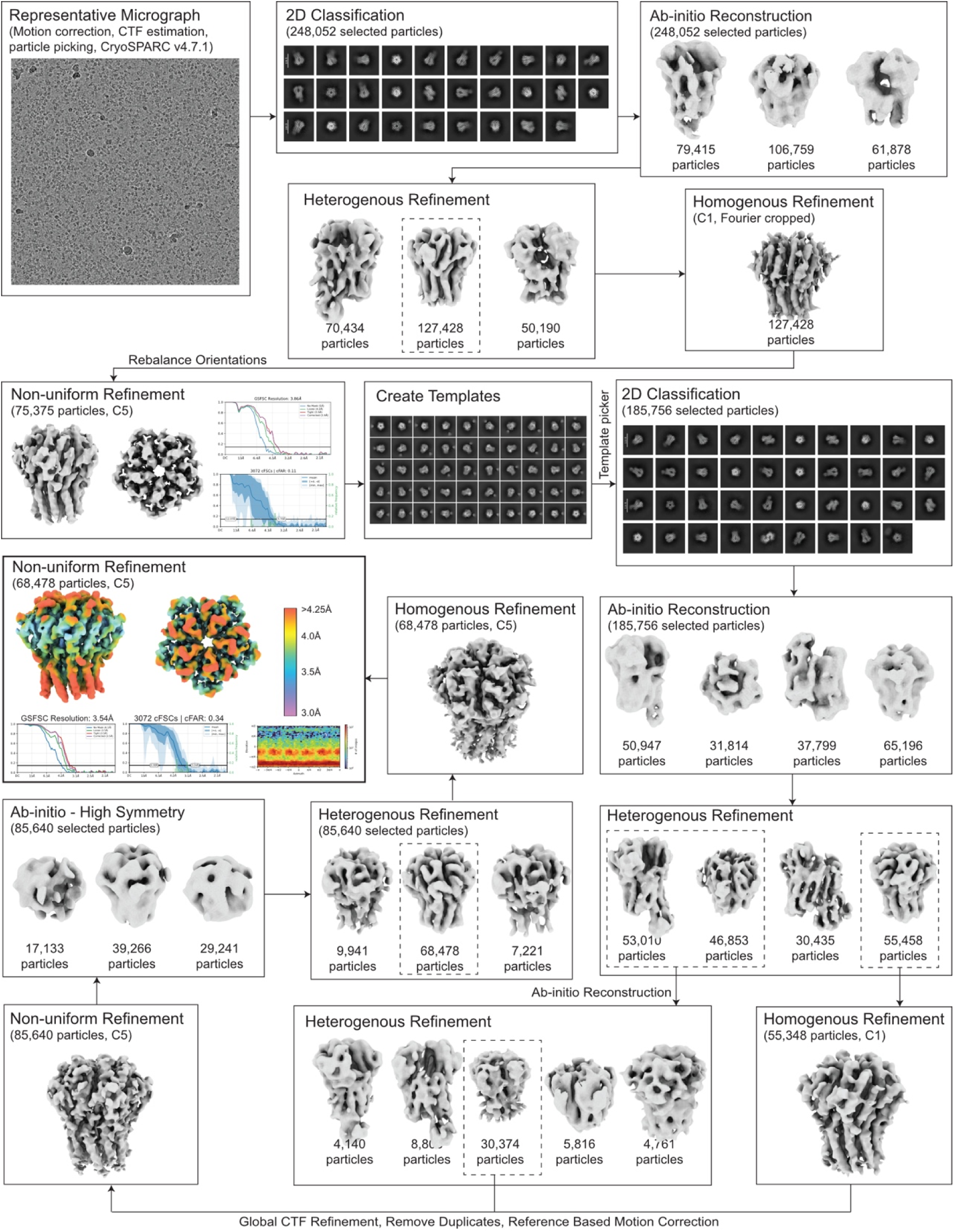
Cryo-EM data processing workflow. Collected micrographs were subjected to motion correction, CTF estimation, iterative particle picking, *ab initio* reconstruction, heterogeneous refinement, homogeneous refinement, global CTF refinement, reference-based motion correction, and non-uniform refinement in cryoSPARC v4.7.1^53^. The final reconstruction was refined with C5 symmetry applied and sharpened using EMReady2^54^.

## Notes

https://github.com/ProteinDesignLab/caliby

### Competing Interest Statement

The authors have declared no competing interest.

### Summary of Updates

Updated to include wet lab validation and additional computational experiments

## References

1. Ingraham, J., Garg, V., Barzilay, R. & Jaakkola, T. Generative models for graph-based protein design. Adv. Neural Inf. Process. Syst. 32, (2019).

2. Hsu, C. et al. Learning inverse folding from millions of predicted structures. In International conference on machine learning 8946–8970 (PMLR, 2022).

3. Dauparas, J. et al. Robust deep learning–based protein sequence design using ProteinMPNN. Science 378, 49–56 (2022).

4. Ingraham, J. B. et al. Illuminating protein space with a programmable generative model. Nature 623, 1070–1078 (2023).

5. Dauparas, J. et al. Atomic context-conditioned protein sequence design using LigandMPNN. Biorxiv 2023–12 (2023).

6. Ruffolo, J. A. et al. Adapting protein language models for structure-conditioned design. bioRxiv 2024–08 (2024).

7. Widatalla, T., Shuai, R. W., Hie, B. & Huang, P. Sidechain conditioning and modeling for full-atom protein sequence design with FAMPNN. in Forty-second International Conference on Machine Learning.

8. Birnbaum, F. & Keating, A. E. Beyond native sequence recovery: Improved modeling of the sequence-energy landscape of protein structures. Proc. Natl. Acad. Sci. 123, e2535494123 (2026).

9. Lu, T., Liu, M., Chen, Y., Kim, J. & Huang, P.-S. Assessing generative model coverage of protein structures with SHAPES. bioRxiv (2025).

10. Onuchic, J. N., Luthey-Schulten, Z. & Wolynes, P. G. Theory of protein folding: the energy landscape perspective. Annu. Rev. Phys. Chem. 48, 545–600 (1997).

11. Ferreiro, D. U., Hegler, J. A., Komives, E. A. & Wolynes, P. G. Localizing frustration in native proteins and protein assemblies. Proc. Natl. Acad. Sci. 104, 19819–19824 (2007).

12. Jenik, M. et al. Protein frustratometer: a tool to localize energetic frustration in protein molecules. Nucleic Acids Res. 40, W348–W351 (2012).

13. Lu, T. et al. Conditional Protein Structure Generation with Protpardelle-1c. bioRxiv 2025–08 (2025).

14. Jumper, J. et al. Highly accurate protein structure prediction with AlphaFold. nature 596, 583–589 (2021).

15. Li, A. J. et al. Neural network-derived Potts models for structure-based protein design using backbone atomic coordinates and tertiary motifs. Protein Sci. 32, e4554 (2023).

16. Sun, H., Dai, H., Dai, B., Zhou, H. & Schuurmans, D. Discrete langevin samplers via wasserstein gradient flow. in International Conference on Artificial Intelligence and Statistics 6290–6313 (PMLR, 2023).

17. Berman, H. M. et al. The Protein Data Bank. Nucleic Acids Res. 28, 235–242 (2000).

18. Hinton, G. E. Products of experts. 9th International Conference on Artificial Neural Networks: ICANN ‘99 vol. 1999 1–6 (1999).

19. Butcher, J., et al. De novo design of all-atom biomolecular interactions with rfdiffusion3. bioRxiv (2025).

20. Lee, J. S. & Kim, P. M. FlowPacker: Protein side-chain packing with torsional flow matching. bioRxiv 2024–07 (2024).

21. Goverde, C. A. et al. Computational design of soluble and functional membrane protein analogues. Nature 631, 449–458 (2024).

22. Pacesa, M. et al. One-shot design of functional protein binders with BindCraft. Nature 646, 483–492 (2025).

23. Morcos, F., et al. Direct-coupling analysis of residue coevolution captures native contacts across many protein families. Proc. Natl. Acad. Sci. 108, E1293–E1301 (2011).

24. Wang, H. et al. Disentanglement of Evolutionary Constraints in Statistical Models of Proteins. PRX Life 2, 023005 (2024).

25. Akiyama, Y. et al. Expanding the scope of protein language modeling to protein-protein interactions with MSA Pairformer. Cell (2026).

26. Zhang, Z., et al. Protein language models learn evolutionary statistics of interacting sequence motifs. Proc. Natl. Acad. Sci. 121, e2406285121 (2024).

27. Lin, Z. et al. Evolutionary-scale prediction of atomic-level protein structure with a language model. Science 379, 1123–1130 (2023).

28. Davtyan, A. et al. AWSEM-MD: protein structure prediction using coarse-grained physical potentials and bioinformatically based local structure biasing. J. Phys. Chem. B 116, 8494–8503 (2012).

29. Watson, J. L. et al. De novo design of protein structure and function with RFdiffusion. Nature 620, 1089–1100 (2023).

30. Wang, K. et al. The structure of phosphatidylinositol remodeling MBOAT7 reveals its catalytic mechanism and enables inhibitor identification. Nat. Commun. 14, 3533 (2023).

31. Rajan, A. et al. BPP43_05035 is a Brachyspira pilosicoli cell surface adhesin that weakens the integrity of the epithelial barrier during infection. Gut Microbes 16, 2409247 (2024).

32. Mattocks, J. A. et al. Enhanced rare-earth separation with a metal-sensitive lanmodulin dimer. Nature 618, 87–93 (2023).

33. Cotruvo Jr, J. A., Featherston, E. R., Mattocks, J. A., Ho, J. V. & Laremore, T. N. Lanmodulin: a highly selective lanthanide-binding protein from a lanthanide-utilizing bacterium. J. Am. Chem. Soc. 140, 15056–15061 (2018).

34. Catoiu, E. A. et al. QSProteome: a community-driven interactive platform for large-scale exploration and evaluation of predicted protein complex structures. Nucleic Acids Res. 54, D661–D672 (2026).

35. Evans, R., et al. Protein complex prediction with AlphaFold-Multimer. bioRxiv 10.1101/2021.10.04.463034 (2022) doi:10.1101/2021.10.04.463034.

36. El Nesr, G., et al. Zero-shot design of a de novo metalloenzyme. bioRxiv 2026–04 (2026).

37. Claude Science & Shanehsazzadeh, A. Autonomous de Novo Protein Binder Design with Claude. https://www-cdn.anthropic.com/30bf50e22a01388bb29bf077ee3f244531594b7a.pdf (2026).

38. Gordon, C., Lu, A. X. & Abbeel, P. Protein language model fitness is a matter of preference. bioRxiv 2024–10 (2024).

39. Pugh, C. W., Nuñez-Valencia, P. G., Dias, M. & Frazer, J. From Likelihood to Fitness: Improving Variant Effect Prediction in Protein and Genome Language Models. bioRxiv 2025–05 (2025).

40. Davey, J. A. & Chica, R. A. Multistate approaches in computational protein design. Protein Sci. 21, 1241–1252 (2012).

41. Praetorius, F. et al. Design of stimulus-responsive two-state hinge proteins. Science 381, 754–760 (2023).

42. Guo, A. B. et al. Deep learning–guided design of dynamic proteins. Science 388, eadr7094 (2025).

43. Dobson, L. et al. UniTmp: unified resources for transmembrane proteins. Nucleic Acids Res. 52, D572–D578 (2024).

44. Karras, T., Aittala, M., Aila, T. & Laine, S. Elucidating the design space of diffusion-based generative models. Adv. Neural Inf. Process. Syst. 35, 26565–26577 (2022).

45. Wohlwend, J. et al. Boltz-1 democratizing biomolecular interaction modeling. BioRxiv 2024–11 (2025).

46. Steinegger, M. & Söding, J. MMseqs2 enables sensitive protein sequence searching for the analysis of massive data sets. Nat. Biotechnol. 35, 1026–1028 (2017).

47. Corley, N. et al. Accelerating Biomolecular Modeling with AtomWorks and RF3. bioRxiv 2025–08 (2025).

48. Ovchinnikov, S., et al. sokrypton/ColabDesign. Zenodo 10.5281/zenodo.13309080 (2025).

49. Mirdita, M. et al. ColabFold: making protein folding accessible to all. Nat. Methods 19, 679–682 (2022).

50. Seemayer, S., Gruber, M. & Söding, J. CCMpred—fast and precise prediction of protein residue–residue contacts from correlated mutations. Bioinformatics 30, 3128–3130 (2014).

51. Passaro, S., et al. Boltz-2: Towards accurate and efficient binding affinity prediction. BioRxiv (2025).

52. Liu, Y. et al. Single-point mutated lanmodulin as a high-performance MRI contrast agent for vascular and kidney imaging. Nat. Commun. 15, 9834 (2024).

53. Punjani, A., Rubinstein, J. L., Fleet, D. J. & Brubaker, M. A. cryoSPARC: algorithms for rapid unsupervised cryo-EM structure determination. Nat. Methods 14, 290–296 (2017).

54. Cao, H. et al. EMReady2: improvement of cryo-EM and cryo-ET maps by local quality-aware deep learning with Mamba. Nat. Commun. (2026).

55. Emsley, P., Lohkamp, B., Scott, W. G. & Cowtan, K. Features and development of Coot. Biol. Crystallogr. 66, 486–501 (2010).

56. Croll, T. I. ISOLDE: a physically realistic environment for model building into low-resolution electron-density maps. Biol. Crystallogr. 74, 519–530 (2018).

57. Pettersen, E. F. et al. UCSF ChimeraX: Structure visualization for researchers, educators, and developers. Protein Sci. 30, 70–82 (2021).

58. Afonine, P. V. et al. Real-space refinement in PHENIX for cryo-EM and crystallography. Biol. Crystallogr. 74, 531–544 (2018).

