## Supplementary information for "Isolation of structural constraints by ensemble-conditioned protein sequence design"

**Supplementary Note 1**

Here we provided additional details for the ensemble-conditioned Potts model formulation used by Caliby.

**Ensemble design as an average of Potts models**

By linearity, the ensemble-conditioned energy $E_{\text{ens}}$ is itself a Potts energy computed from Potts parameters equal to the average of the per-structure parameters:

$E_{\text{ens}}\left( S \right)$

$=\frac{1}{K}\sum_{k=1}^{K} E_{k}\left( S \right)$

$=\frac{1}{K}\sum_{k=1}^{K} \left( \sum_{i} h_{i}^{\left( k \right)}\left( s_{i} \right)+\sum_{i<j} J_{ij}^{\left( k \right)}\left( s_{i},s_{j} \right) \right)$

$=\frac{1}{K}\left( \sum_{i} \sum_{k=1}^{K} h_{i}^{\left( k \right)}\left( s_{i} \right)+\sum_{i<j} \sum_{k=1}^{K} J_{ij}^{\left( k \right)}\left( s_{i},s_{j} \right) \right)\quad\text{(swap sums)}$

$=\sum_{i} \left( \frac{1}{K}\sum_{k=1}^{K} h_{i}^{\left( k \right)} \right)\left( s_{i} \right)+\sum_{i<j} \left( \frac{1}{K}\sum_{k=1}^{K} J_{ij}^{\left( k \right)} \right)\left( s_{i},s_{j} \right)$

As a result, by simply averaging the Potts parameters across the input structures, we can repeatedly sample sequences for the ensemble at the same rate as sampling for a single structure. This makes Caliby scale more computationally efficiently with ensemble size $K$. Given a protein of length $N$, ProteinMPNN’s autoregressive decoding requires $K\times N$ forward passes, incurring additional cost for each sampled sequence. By contrast, Caliby precomputes Potts parameters with one forward pass per conformer and samples sequences with cost independent of $K$ thereafter.

Since precomputing the Potts parameters for each structure requires only a single pass through the neural network, the sampling time is dominated by sequence sampling from the Potts model. In practice, on a backbone of length 500, computing Potts parameters takes roughly 0.05 seconds per input structure and 0.8 seconds to sample each sequence from the averaged Potts model on an NVIDIA H100.

**Ensemble-conditioned sampling as a normalized geometric mean**

Ensemble-conditioned Caliby can be seen as sampling sequences from a normalized geometric mean over the per-structure sequence distributions. If the ensemble energy is the average $E_{\text{ens}}\left( S \right)=\frac{1}{K}\sum_{k=1}^{K} E_{k}\left( S \right)$, then

$$p_{\text{ens}}\left( S \right)\propto\exp\left( -\frac{1}{K}\sum_{k=1}^{K} E_{k}\left( S \right) \right)=\left( \prod_{k=1}^{K} e^{-E_{k}\left( S \right)} \right)^{1/K}\propto\prod_{k=1}^{K} p_{k}\left( S \right)^{1/K}$$

So the ensemble Potts distribution is proportional to the geometric mean of the per-structure Potts distributions.

**ProteinMPNN tied sampling as a per-step normalized geometric mean**

At each decoding step, ProteinMPNN tied sampling over an ensemble is a normalized geometric mean of the per-structure distributions for the next token. However, we note that unlike with ensemble-conditioned Caliby, this does not necessarily hold for the whole sequence distribution. To see this, at decoding step $t$, for a given amino acid $a$, structure $k$ yields logits $z_{a}^{\left( k \right)}$ with

$$p^{\left( k \right)}\left( a \right)=\frac{e^{z_{a}^{\left( k \right)}}}{Z^{\left( k \right)}},\quad\quad Z^{\left( k \right)}=\sum_{b} e^{z_{b}^{\left( k \right)}}$$

The tied distribution at decoding step $t$ is obtained by averaging logits, then applying softmax:

$p_{\text{ens}}\left( a \right)=\text{softmax}\left( \frac{1}{K}\sum_{k} z_{a}^{\left( k \right)} \right)$

$=\frac{e^{\frac{1}{K}\sum_{k} z_{a}^{\left( k \right)}}}{\sum_{b} e^{\frac{1}{K}\sum_{k} z_{b}^{\left( k \right)}}}$

$=\frac{\left( \prod_{k} e^{z_{a}^{\left( k \right)}} \right)^{1/K}}{\sum_{b} \left( \prod_{k} e^{z_{b}^{\left( k \right)}} \right)^{1/K}}$

From the previous line, the denominator is independent of the choice of amino acid $a$ and can be treated as a normalization constant, so we can see that

$$p_{\text{ens}}\left( a \right)\propto\left( \prod_{k} p^{\left( k \right)}\left( a \right) \right)^{1/K}$$

meaning that at each decoding step, the tied sampling distribution is a normalized geometric mean over the next-token distributions predicted from each structure.

In practice, for ProteinMPNN tied sampling, we used the implementation provided by the Kuhlman Lab at <https://github.com/Kuhlman-Lab/proteinmpnn>.

**Supplementary Figures and Tables**

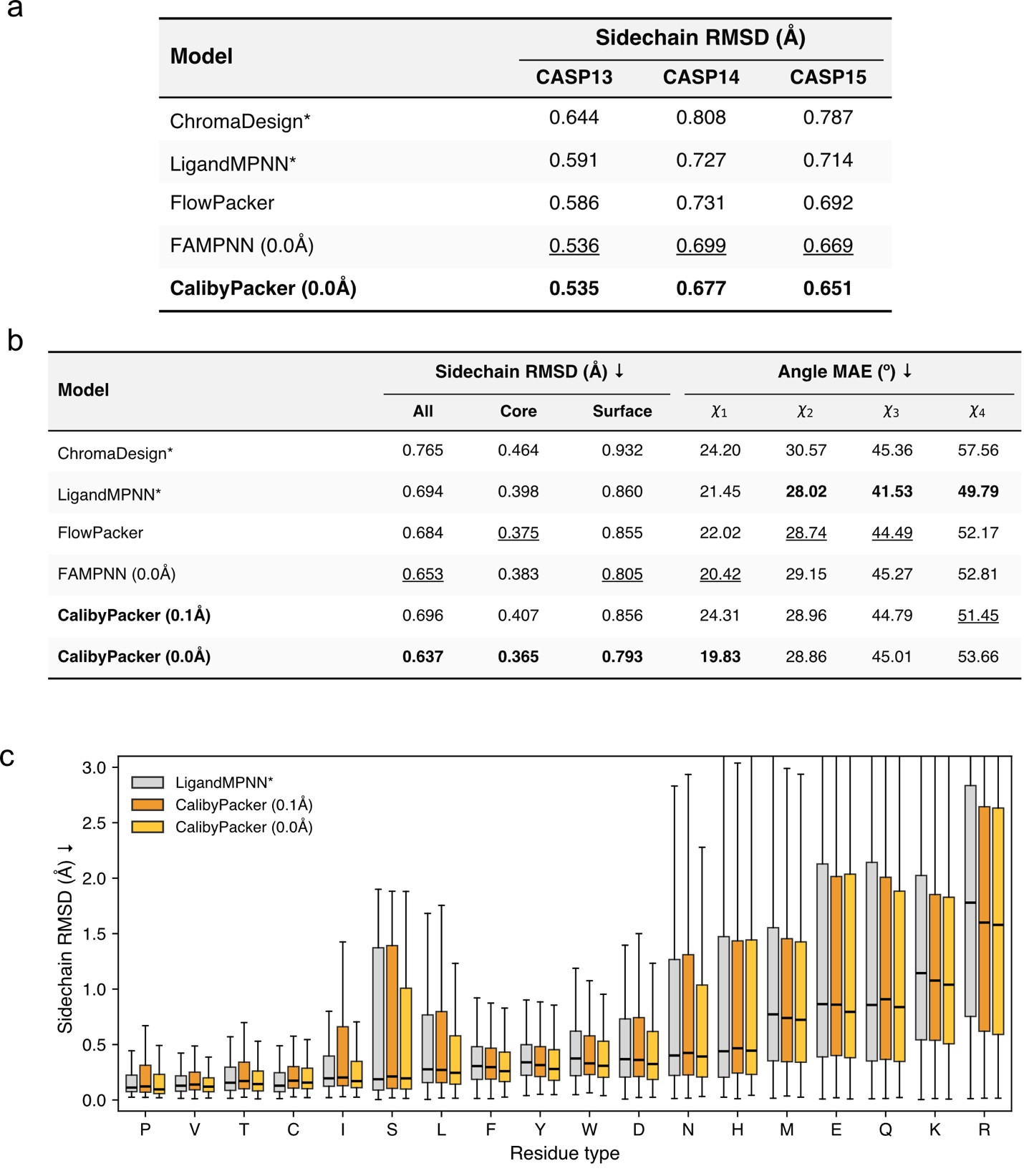

**Supplementary Figure 1. Sidechain packing evaluation. (a)** Sidechain packing performance comparison across CASP datasets. **(b)** Sidechain RMSD and dihedral angle MAE across CASP13, 14, and 15 targets (n = 99) held out at 40% sequence similarity. **(c)** Per-residue sidechain RMSD for LigandMPNN, CalibyPacker (0.1Å), and CalibyPacker (0.0Å). For (a) and (b), bold indicates best performance; underline indicates second best. Methods marked with an asterisk were trained on datasets that do not explicitly hold out CASP proteins. RMSD for all methods are computed accounting for sidechain symmetry in Asp, Glu, Phe, and Tyr and may differ from previously reported values.

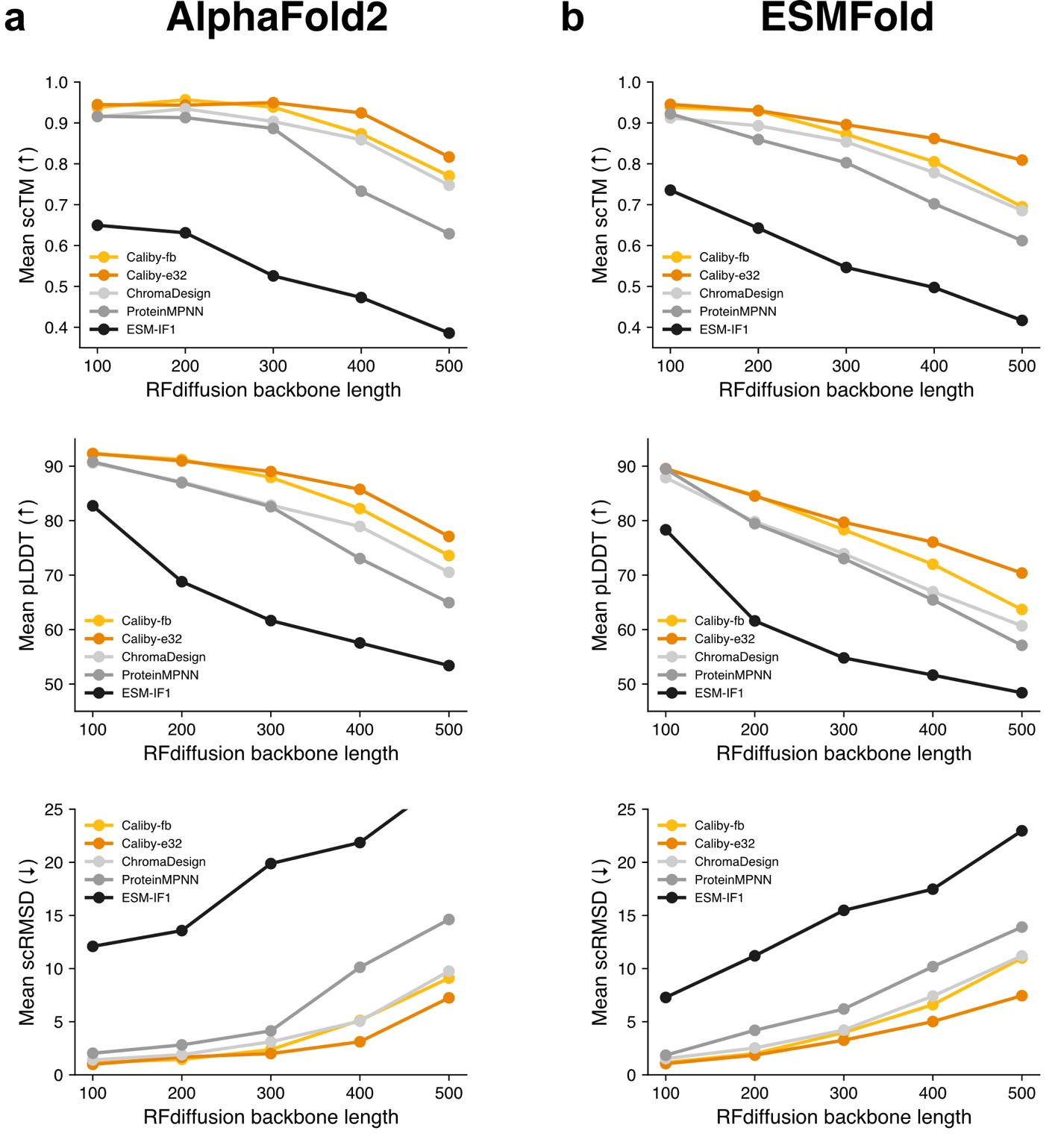

**Supplementary Figure 2. RFdiffusion backbone designability across sequence design methods.** **(a)** Mean single-sequence AlphaFold2 designability metrics (scTM, pLDDT, scRMSD) as a function of RFdiffusion backbone length for Caliby-fb, Caliby-e32, ChromaDesign, ProteinMPNN, and ESM-IF1 (100 backbones per length). **(b)** Same as (a) but with ESMFold instead of single-sequence AlphaFold2.

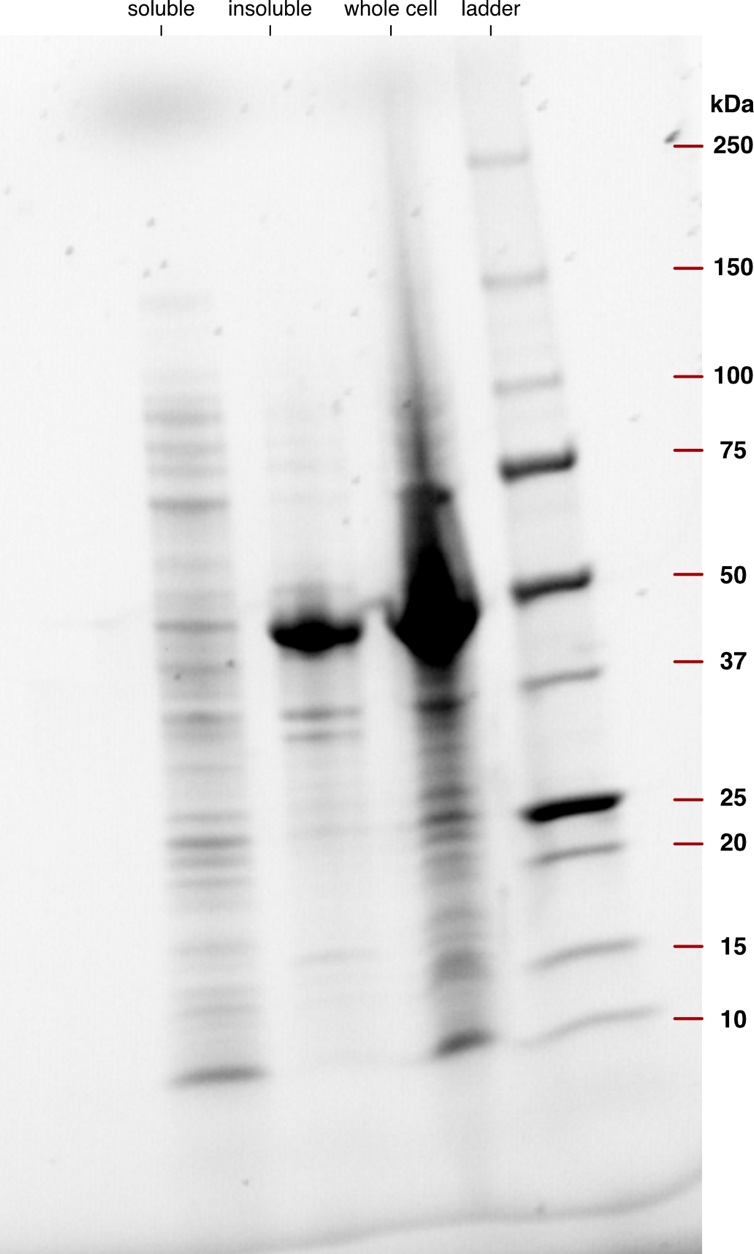

**Supplementary Figure 3. The SolubleMPNN design for 8ERC is found predominantly in the insoluble fraction.** SDS-PAGE of soluble, insoluble, and whole-cel fractions from *E. coli* expressing the SolubleMPNN design for membrane-bound O-acyltransferase 8ERC. A band at the expected mass of the design (53 kDa) appears in the whole-cell and insoluble fractions but not the soluble fraction.

| **EM data collection** | **Caliby designed Q8XAZ3** |
| --- | --- |
| Microscope | Glacios |
| Voltage (kV) | 200 |
| Detector | Falcon 4i |
| Magnification (nominal) | 150,000 × |
| Pixel size (Å/pix) | 0.93 |
| Exposure (e^-^/Å^2^) | 40.1 |
| Defocus range (μm) | 0.9–2.7 |
| Automation software | EPU |
| Sample | Caliby designed Q8XAZ3 at 6 mg/mL |
| Tilt angle ( ̊ ) | 0 |
| Micrographs collected | 728 |
| Micrographs used | 707 |
| Particles extracted (total) | 691,997 |
| **3D reconstruction statistics** |  |
| Particles | 68,478 |
| Symmetry | C5 |
| Map sharpening B-factor | –143.3 |
| Unmasked resolution at 0.5 FSC (Å) | 6.7 |
| Masked resolution at 0.5 FSC (Å) | 4.0 |
| Unmasked resolution at 0.143 FSC (Å) | 5.4 |
| Masked resolution at 0.143 FSC (Å) | 3.5 |
| **Model refinement and validation statistics** |  |
| Refinement package | ChimeraX ISOLDE, Phenix, Coot |
| Refinement tool | Phenix real-space refinement |
| Refinement strategies | min global, local_grid_search, adp reference model restraints, ss restraints, rotamer restraints, Ramachandran restraints |
| Composition |  |
| Amino acids | 1,070 |
| RMSD bonds (Å) | 0.005 |
| RMSD angles (º) | 0.994 |
| Average B-factors |  |
| Amino acids | 91.9 |
| Ramachandran |  |
| Favored (%) | 98.1 |
| Allowed (%) | 1.9 |
| Outliers (%) | 0 |
| Rotamer outliers (%) | 0 |
| Clash score | 5.31 |
| C-beta outliers (%) | 0 |
| CaBLAM outliers (%) | 1.46 |
| CC (mask) | 0.85 |
| MolProbity score | 1.28 |
| EMRinger score | 1.15 |
| PDB ID | 38DW |
| EMDB ID | EMD-78753 |

**Supplementary Table 1. Cryo-EM data collection and model building statistics.**

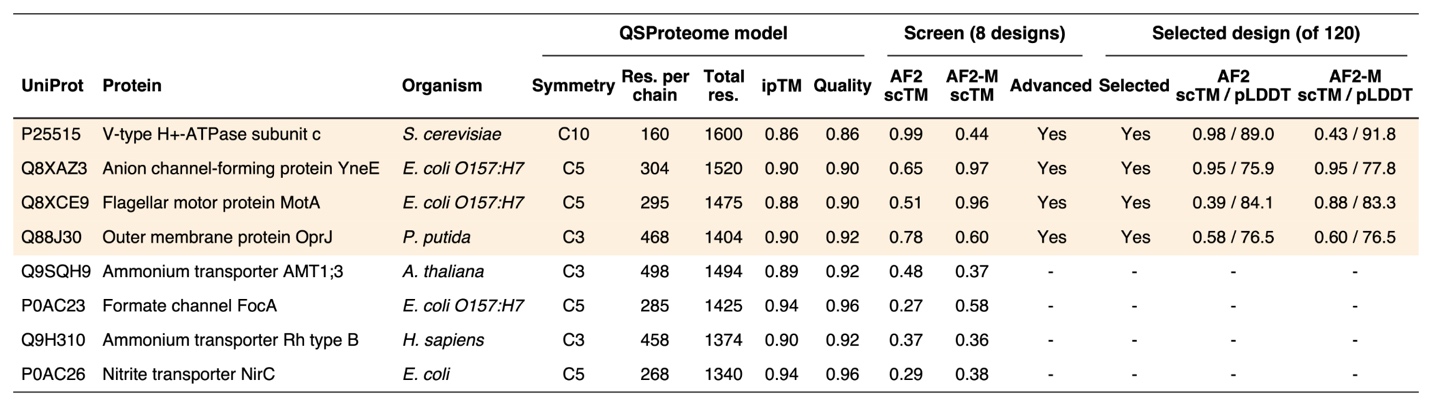

**Supplementary Table 2. Homo-oligomeric membrane protein assemblies selected from QSProteome and design outcomes.** Selected criteria and design procedure as described in Methods. Shaded rows advanced past the initial 8-design screen, while dashes indicate targets that did not advance.

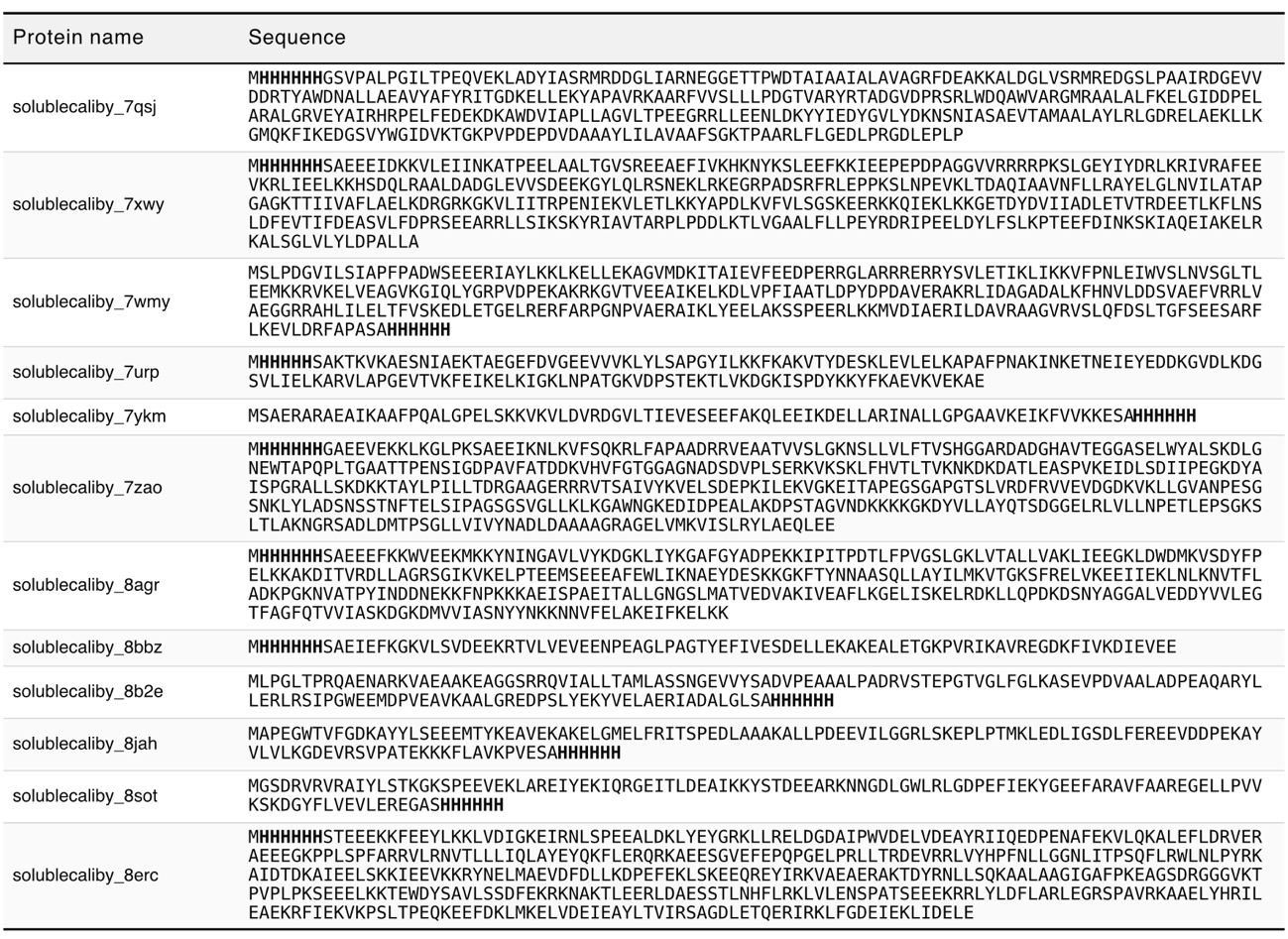

**Supplementary Table 3. SolubleCaliby-e32 sequences for native protein redesigns.** His-tags are shown in bold.

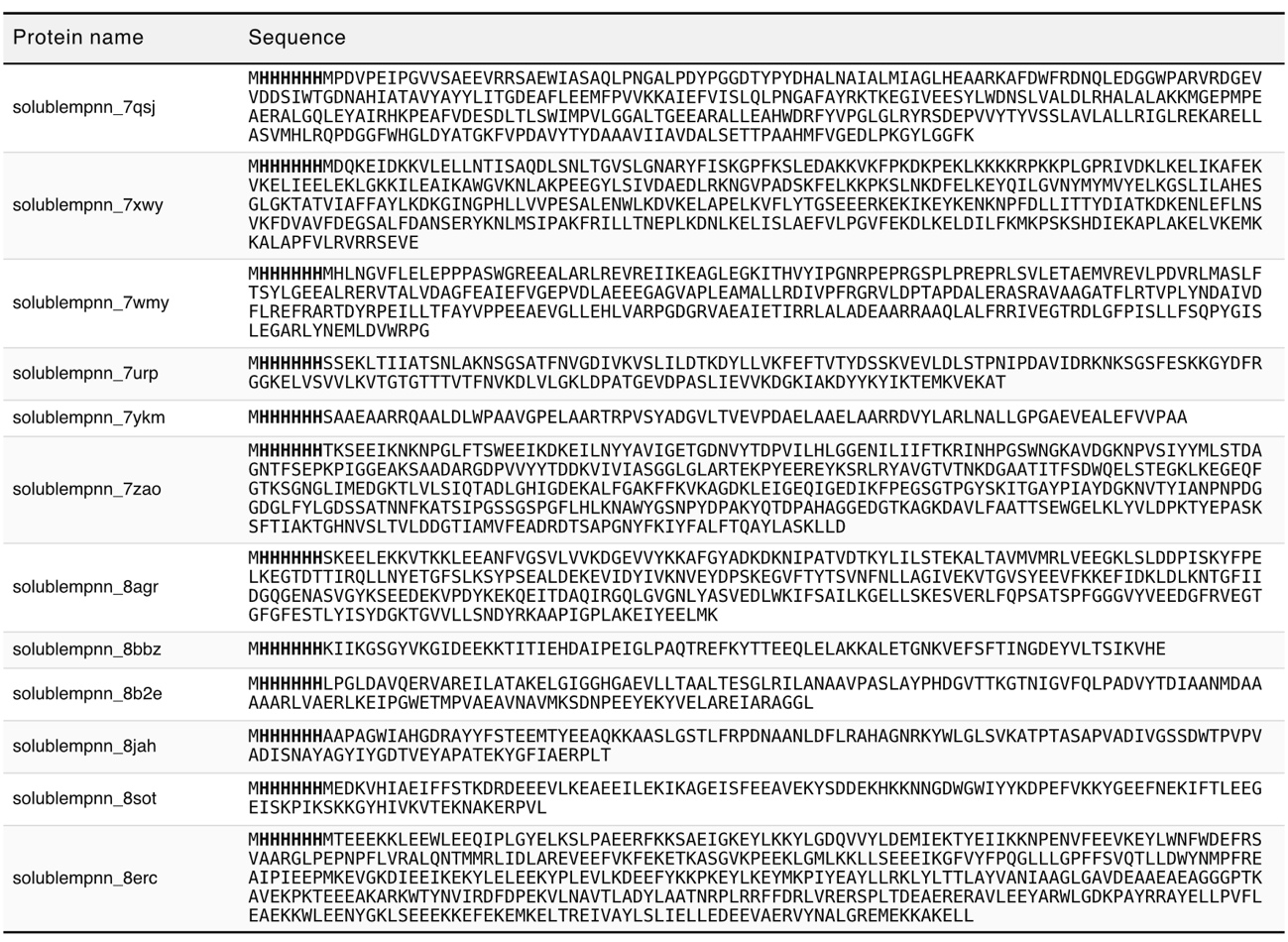

**Supplementary Table 4. SolubleMPNN sequences for native protein redesigns**. His-tags are shown in bold.
